# Ca^2+^-dependent mechanism of membrane insertion and destabilization by the SARS-CoV-2 fusion peptide

**DOI:** 10.1101/2020.12.03.410472

**Authors:** George Khelashvili, Ambrose Plante, Milka Doktorova, Harel Weinstein

## Abstract

Cell penetration after recognition of the SARS-CoV-2 virus by the ACE2 receptor, and the fusion of its viral envelope membrane with cellular membranes, are the early steps of infectivity. A region of the Spike protein (S) of the virus, identified as the “fusion peptide” (FP), is liberated at its N-terminal site by a specific cleavage occurring in concert with the interaction of the receptor binding domain of the Spike. Studies have shown that penetration is enhanced by the required binding of Ca^2+^ ions to the FPs of corona viruses, but the mechanisms of membrane insertion and destabilization remain unclear. We have predicted the preferred positions of Ca^2+^ binding to the SARS-CoV-2-FP, the role of Ca^2+^ ions in mediating peptide-membrane interactions, the preferred mode of insertion of the Ca^2+^-bound SARS-CoV-2-FP and consequent effects on the lipid bilayer from extensive atomistic molecular dynamics (MD) simulations and trajectory analyses. In a systematic sampling of the interactions of the Ca^2+^-bound peptide models with lipid membranes SARS-CoV-2-FP penetrated the bilayer and disrupted its organization only in two modes involving different structural domains. In one, the hydrophobic residues F833/I834 from the middle region of the peptide are inserted. In the other, more prevalent mode, the penetration involves residues L822/F823 from the LLF motif which is conserved in CoV-2-like viruses, and is achieved by the binding of Ca^2+^ ions to the D830/D839 and E819/D820 residue pairs. FP penetration is shown to modify the molecular organization in specific areas of the bilayer, and the extent of membrane binding of the SARS-CoV-2 FP is significantly reduced in the absence of Ca^2+^ ions. These findings provide novel mechanistic insights regarding the role of Ca^2+^ in mediating SARS-CoV-2 fusion and provide a detailed structural platform to aid the ongoing efforts in rational design of compounds to inhibit SARS-CoV-2 cell entry.

**STATEMENT OF SIGNIFICANCE:** SARS-CoV-2, the cause of the COVID-19 pandemic, penetrates host cell membranes and uses viral-to-cellular membrane fusion to release its genetic material for replication. Experiments had identified a region termed “fusion peptide” (FP) in the Spike proteins of coronaviruses, as the spearhead in these initial processes, and suggested that Ca^2+^ is needed to support both functions. Absent structure and dynamics-based mechanistic information these FP functions could not be targeted for therapeutic interventions. We describe the development and determination of the missing information from analysis of extensive MD simulation trajectories, and propose specific Ca^2+^-dependent mechanisms of SARS-CoV-2-FP membrane insertion and destabilization. These results offer a structure-specific platform to aid the ongoing efforts to use this target for the discovery and/or of inhibitors.

## INTRODUCTION

The corona viruses (CoV) are envelope viruses that infect human and animal cells via fusion of the viral envelope membrane with cellular membranes ^1^. Members of the CoV family include the well-known agents of recent pandemics such as severe acute respiratory syndrome CoV (SARS-CoV), the Middle East respiratory syndrome CoV (MERS-CoV) ^2^, and the severe acute respiratory syndrome CoV-2 (SARS-CoV-2) virus responsible for the currently ongoing CoVID-19 pandemic ^3,4^.

The first point of the connection between the SARS-CoV-2 virus and the human cell is at the ACE2 receptor ^1,5–7^, and is now quite well characterized structurally ^8–17^. It is mediated by the Spike protein which is a multi-domain homo-trimer glycoprotein anchored in the viral envelope membrane ^5,18^. Each monomer of the spike protein consists of two subunits, S1 which includes the receptor binding domain (RBD) that attaches to the ACE2 receptor ^6,7^, and S2 which includes a segment termed “fusion peptide” that is central to the fusion process leading to cell infection by CoVs ^5,19^. A catalytic cleavage site (the S1/S2 site) is recognized by a furin-like protease that can separate the two component domains of the Spike monomer ^20^. Host cell recognition of both SARS-CoV and SARS-CoV-2 involves the angiotensin converting enzyme-2 (ACE2) which binds the RBD on S1. The binding of the S1 domain to the host receptor exposes another cleavage site, the S2’ site on the S2 domain, to processing by cell surface enzymes of the host (primarily TMPRSS2) ^7^. Cleavage of the S2’ site occurs at the N-terminus of the fusion peptide (FP) segment, and a “tectonic” change in the conformation of the S2 domain repositions the FP regions of each of the three monomers close to one another to form a “spearhead” of the Spike, ready to penetrate the cell membrane while still attached to the S1 trimeric bundle ^12,21^.

Fusion Peptide insertion is thought to result in destabilization of the target cell membrane ^22^ as an essential step in allowing the entire virus to penetrate into the cell. In the cell, the FP spearheads the fusion of the viral capsule membrane with cell membranes to release the encapsulated genetic material for replication. Thus far, even the rapid pace of discovery of structural information about the SARS-CoV-2 virus ^8–13,23^, has not yielded molecular level data about the vital step of engagement of the viral Spike protein with the membrane, much less the structure of its spearhead. Even less is known at the needed molecular level about the mechanism by which the FPs in the spearhead are involved in the fusion of the cell and virus membranes, despite the wealth of information that has recently accumulated from molecular dynamics (MD) simulations on SARS-CoV-2 models (e.g., see Refs. ^24–29^).

Here we present results from an atomistic-level MD investigation of the interactions of SARS-CoV-2 FPs with membranes, and the consequences of these interactions on membrane properties. Biophysical and structure/function studies of the SARS-CoV-2 FP ^30^ as well as of the FPs from SARS-CoV ^31^ and MERS-CoV ^32^, which are close homologs of SARS-CoV-2, had produced important insights about the functional significance of specific regions of the FP such as the large number of conserved acidic residues in the two consecutive fragments of the FP termed FP1 and FP2 (residues 816-835 and 835-854, respectively, in SARS-CoV-2), and the LLF hydrophobic motif ^19,31,33^ (L821/L822/F823 in SARS-CoV-2, see Figure 1). Studies on SARS-CoV-2 FP ^30^ and SARS-CoV ^31^ have suggested that together, the F1 and F2 fragments form a bipartite membrane interaction platform, and that acidic residues in both FP1 and FP2 promote membrane binding through their interactions with Ca^2+^ ions ^30,31^. These findings about the functional role of Ca^2+^ are echoed by results obtained for FPs from other related viruses, such as MERS-CoV ^32^ and Ebola virus ^34^. Indeed, in the MERS-CoV FP one of these acidic residues, E891 in the N-terminal (FP1) part (corresponding to E819 in SARS-CoV-2 numbering, Figure 1A) emerged as crucial for Ca^2+^ interactions and fusion-related membrane perturbation effects ^32^. In contrast to the SARS-CoV FP and SARS-CoV-2 FP, however, isothermal titration calorimetry (ITC) experiments with the MERS-CoV FP showed that the latter binds only a single Ca^2+^ ion ^32^. Notably, much of the structural and functional inferences obtained from these studies appears to be in excellent agreement with, and recapitulated, by the recently published results obtained from equivalent experiments with the SARS-CoV-2 FP ^30^.

**Figure 1:**
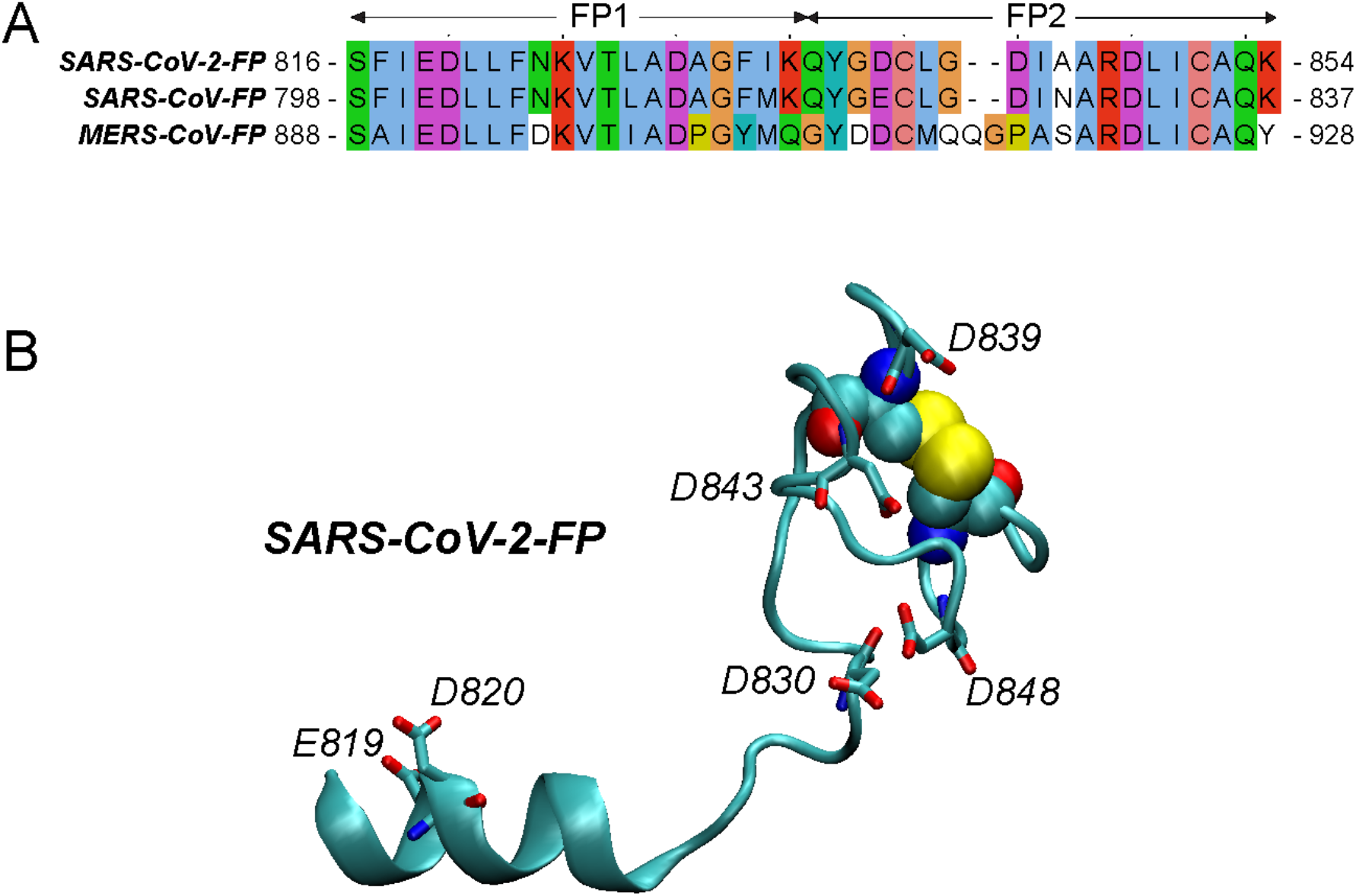
(**A**) Sequence alignment of the fusion peptide (FP) segments from the spike proteins of SARS-CoV-2 (top), SARS-CoV (middle), and MERS-CoV (bottom) coronaviruses. The sequences are colored according to ClustalX scheme. The conserved acidic residues are shown in purple, and the conserved Cys residues, forming a disulfide bond (C840 and C851 in SARS-CoV-2-FP numbering), are highlight in orange. FP1 and FP2 fragments of the peptide are marked and labeled. (**B**) Structural model of SARS-CoV-2-FP used in this study. The acidic residues are shown in licorice and labeled. The Cys residues forming the disulfide bond are shown in space fill.

The involvement of the conserved LLF motif in the FP1 fragment of the SARS-CoV-FP as a critical structural element for membrane insertion and fusogenicity of the peptide was identified from mutagenesis studies ^19,31,33^. Also, perturbation of the lipid bilayer by Ca^2+^-dependent FP-membrane interactions was shown with Electron Spin Resonance (ESR) experiments to affect the organization of the lipid head group and backbone atoms of the lipids at the interaction site, but not the central region of the membrane hydrophobic core ^30,31, 32,34^. Still, the loci of Ca^2+^ binding to CoV FPs and their role in any specific (but unknown) modes of FP interaction with the membrane, had remained undetermined. This has hindered the interpretation of any measurable effects of viral interactions with the membrane, and thus any approaches to mitigate infectivity by targeting this essential region.

We addressed these fundamental unknowns about the SARS-CoV-2-FPs and their structure-based dynamic mechanisms with extensive atomistic ensemble molecular dynamics (MD) simulations. From analyses of the various trajectories, we predicted multiple ways in which Ca^2+^ ions engage the FP’s acidic residues singly and in pairs. From massive simulations of spontaneous membrane binding of all feasible variants of Ca^2+^-bound SARS-CoV-2-FP we found two preferred modes of membrane penetration. In one of these modes the inserted hydrophobic residues are F833/I834 from the FP2 region of the peptide which, in the full spike context are closely tethered to the S2 domain. The second more prevalent mode of penetration involves the N-terminal portion of the F1 segment, which is free after the S2’ cleavage. In this mode the hydrophobic insertion consists of the highly conserved residues L822/F823 (LLF motif) and is achieved by the binding of Ca^2+^ ions to the D830/D839 pair and the E819/D820 pair of acidic residues. Notably, our extensive control simulations to evaluate the role of Ca^2+^ binding in the process showed that membrane binding/insertion of the SARS-CoV-2 FP is greatly diminished overall in the absence of Ca^2+^ ions. Moreover, when bound the peptide exerted structural changes in the lipid bilayer whose nature and magnitude showed dependence on the local peptide concentration. Together, these findings provide new mechanistic insights and important molecular-level details about the SARS-CoV-2-FP penetration into the membrane and the structural dynamics underlying the key role of Ca^2+^ binding in this process.

## METHODS

### Molecular dynamics (MD) simulations of the SARS-CoV-2-FP in water

All computations were based on the atomistic structure of the FP (residues 816-854) from the full-length SARS-CoV-2 spike protein model described in Ref. ^27^. In this structure, residues Cys840 and Cys851 form a disulfide bond (see Figure 1). For these studies, the FP segment from the full-length model was isolated and capped with neutral N-and C-termini (ACE and CT3, respectively, in the CHARMM force-field nomenclature). Protonation states of all the titratable residues were predicted at pH 7 using Propka 3.1 software ^35^.

For the atomistic MD simulations of the SARS-CoV-2-FP in water, the peptide was embedded in a rectangular solution box and ionized using VMD tools (“Add Solvation Box” and “Add Ions”, respectively) ^36^. The box of dimensions ~90 Å × 80 Å × 82 Å included an ionic solution mixture of 49 Na^+^, 51 Cl^−^, 2 Ca^2+^ ions, and ~18000 water molecules. The total number of atoms in the system was 54,510.

The system was equilibrated with NAMD version 2.12 ^37^ following a multi-step protocol during which the backbone atoms of the SARS-CoV-2-FP as well as Ca^2+^ ions in the solution were first harmonically constrained and subsequently gradually released in four steps (totaling ~3ns), changing the restrain force constants *k*_F_ from 1, to 0.5, to 0.1 kcal/ (mol Å^2^), and 0 kcal/ (mol Å^2^). These simulations implemented all option for rigidbonds, 1fs (for *k*_F_ 1, 0.5, and 0.1 kcal/ (mol Å^2^)) or 2fs (for *k*_F_ of 0) integration time-step, PME for electrostatics interactions ^38^, and were carried out in NPT ensemble under isotropic pressure coupling conditions, at a temperature of 310 K. The Nose-Hoover Langevin piston algorithm ^39^ was used to control the target P = 1 atm pressure with the “LangevinPistonPeriod” set to 200 fs and “LangevinPistonDecay” set to 50 fs. The van der Waals interactions were calculated applying a cutoff distance of 12 Å and switching the potential from 10 Å.

After this initial equilibration phase, the velocities of all atoms in the system were reset and ensemble MD runs were initiated with OpenMM version 7.4 ^40^ during which the system was simulated in 18 independent replicates, each for 640ns (i.e., cumulative time of ~11.5 μs). These runs implemented PME for electrostatic interactions and were performed at 310K temperature under NVT ensemble. In addition, 4fs time-step was used, with hydrogen mass repartitioning and with “friction” parameter set to 1.0/picosecond. Additional parameters for these runs included: “EwaldErrorTolerance” 0.0005, “rigidwater” True, and “ConstraintTolerance” 0.000001. The van der Waals interactions were calculated applying a cutoff distance of 12 Å and switching the potential from 10 Å.

### Preparation of a pure lipid membrane for atomistic MD simulations

Using the CHARMM-GUI web server ^41^, we designed a symmetric lipid membrane composed of a 3:1:1 mixture of POPC (1-palmitoyl-2-oleoyl-glycero-3-phosphocholine), POPG (1-palmitoyl-2-oleoyl-sn-glycero-3-phospho-(1′-rac-glycerol)), and cholesterol. The total number of lipids in the bilayer was 600. The membrane was solvated and ionized (with 0.15 mM salt concentration), for a total of ~163,000 atoms.

This system was equilibrated with NAMD version 2.12 following the standard multi-step protocol provided by CHARMM-GUI, and then simulated with unbiased MD for 25ns. After this step, three independent replicates were run, each 75ns long. These runs implemented all option for rigidbonds, 2fs integration time-step, PME for electrostatics interactions, and were carried out in NPT ensemble under semi-isotropic pressure coupling conditions, at a temperature of 298 K. The Nose-Hoover Langevin piston algorithm was used to control the target P = 1 atm pressure with the “LangevinPistonPeriod” set to 50 fs and “LangevinPistonDecay” set to 25 fs. The van der Waals interactions were calculated with a cutoff distance of 12 Å and switching the potential from 10 Å. In addition, “vdwForceSwitching” was set to *yes*.

### MD simulations of SARS-CoV-2-FP interactions with lipid membranes

Interactions of a single, or multiple, FPs with lipid membranes were started by placing 1, 3, 6, 9, or 12 replicas of the Ca^2+^-bound SARS-CoV-2-FPs in the proximity of a bilayer composed of 3:1:1 POPC/POPG/Cholesterol that had been pre-equilibrated for 25ns as described above. Parallel simulations of a single SARS-CoV-2-FP in the absence of Ca^2+^ ions were carried out with the same pre-equilibrated 3:1:1 POPC/POPG/Cholesterol bilayer. The starting structure for the FP for the latter runs was the FP model from Ref. ^27^ (see Introduction). The SARS-CoV-2-FP and membrane were embedded in a solution box containing 150 mM Na^+^Cl^−^ salt concentration. The total number of atoms ranged from 167,000 to 217,000, depending on the number of SARS-CoV-2-FPs in the simulation box.

Each system was equilibrated with NAMD version 2.12 following the same multi-step protocol described above during which the backbone atoms of the FP as well as the Ca^2+^ ions were first harmonically constrained and subsequently gradually released in four steps. After this phase, the velocities of all atoms of the system were reset, and ensemble MD runs were initiated with OpenMM version 7.4. Table 1 lists the number of replicates and simulation times for each construct. These runs implemented PME for electrostatic interactions and were performed at 298K temperature under NPT ensemble using semi-isotropic pressure coupling, with 4fs time-steps, using hydrogen mass repartitioning and with “friction” parameter set to 1.0/picosecond. Additional parameters for these runs included: “EwaldErrorTolerance” 0.0005, “rigidwater” True, and “ConstraintTolerance” 0.000001. The van der Waals interactions were calculated applying a cutoff distance of 12 Å and switching the potential from 10 Å.

**Table 1:**
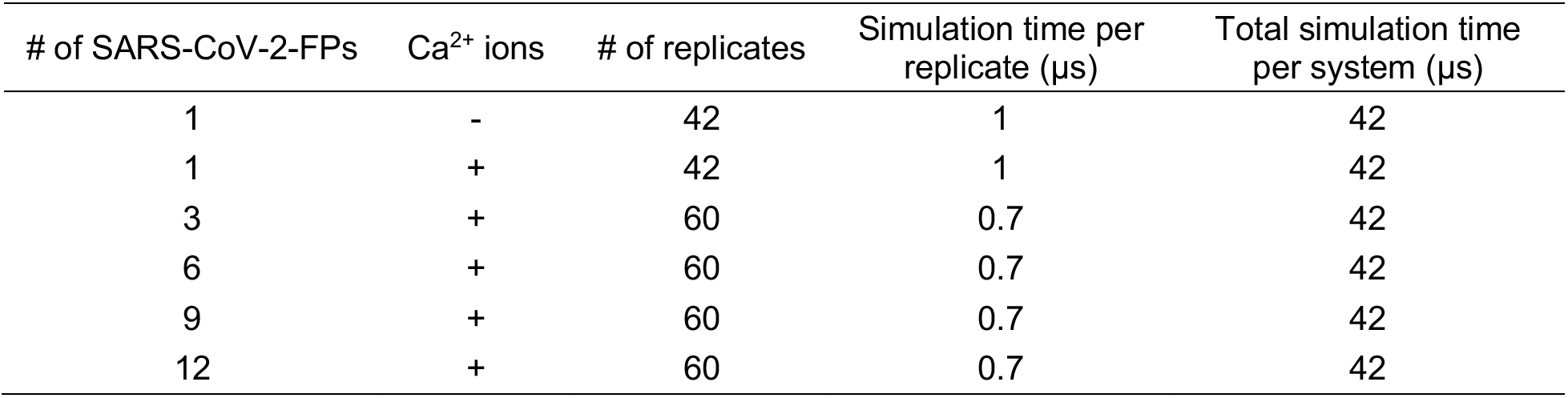
Listing of SARS-CoV-2-FP/lipid membrane systems studied with all-atom ensemble MD simulations. Columns from left to right show: number of SARS-CoV-2-FPs per system; presence or absence of Ca^2+^ ions; number of independent replicates for each construct; simulation time per single replicate; total simulation time per system. The membrane was 3:1:1 mixture of POPC/POPG/Cholesterol, with a total of 600 lipids (see Methods).

For all simulations we used the latest CHARMM36 force-field for proteins and lipids ^42^, as well as the recently revised CHARMM36 force-field for ions which includes non-bonded fix (NBFIX) parameters for Na^+^ and Ca^2+^ ions ^43^.

### Calculation of pressure profiles from MD simulations

Lateral pressure profiles were calculated for selected FP-membrane systems (see Results) as well as for the pure lipid bilayer system described above. For the latter, the last halves of the trajectories of the three 75ns-long independent replicates were combined into a single trajectory for the pressure analysis. Similarly, for the FP-membrane complexes three 75ns long MD trajectory replicas were accumulated in NAMD and the last halves of these trajectories were concatenated and used for analysis. These simulations used the CHARMM36 force field, were initiated from the corresponding last frames of the OpenMM trajectories and were performed with the parameters used in the last step of the multi-stage equilibration protocol described above.

The pressure profiles were calculated with NAMD following the protocol described in Ref. ^44^. The concatenated trajectory for analysis for each system comprised ~26,000 frames with output frequency of 2 ps. The trajectory was first centered on the average z position of the terminal methyl carbons of the lipid chains set at (x,y,z) = (0,0,0). The bilayer was then divided into a discrete number of equal-thickness (~0.8 A) slabs in the direction normal to the bilayer plane. The kinetic, bonded and non-bonded contributions to the pressure were obtained separately from the Ewald contributions, and the two were added to produce the lateral components of the pressure tensor (*p*_*xx*_, *p*_*yy*_) in each slab. The normal component *p_zz_* was calculated as described earlier (see SI in Ref. ^44^) with the formula:

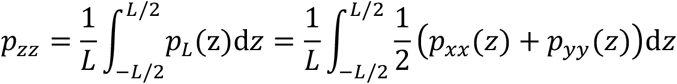

where L is the length of the simulation box in the z direction, and *p_L_*(*z*) represents the tangential component of the pressure tensor. From here, the pressure in each slab was *p*(*z*) = *p_L_*(*z*) − *p*_zz_.

## RESULTS

The FP of the SARS-CoV-2 spike protein (SARS-CoV-2-FP) harbors 6 acidic residues (E819, D820, D830, D839, D843, and D848) that are conserved in many coronaviruses, such as SARS-CoV and MERS-CoV (see Figure 1). As the binding of Ca^2+^ ions was found experimentally to modulate the ability of the FPs from these CoVs to insert into lipid membranes, all these residues had to be considered putative Ca^2+^-binding sites ^31,32,34 30^. To identify the modes of Ca^2+^ interaction with the SARS-CoV-2-FP and to elucidate the function-related consequences of these interactions, we first used atomistic MD simulations of SARS-CoV-2-FP in aqueous solution to explore the spontaneous binding of Ca^2+^ and its effect on the conformation of the peptide (see Methods).

### Modes of Ca^2+^ binding to SARS-CoV-2-FP

Analysis of 18 independent 640ns MD trajectories amounting to >11.5μs sampling (Methods), captured multiple events of association of Ca^2+^ ions with the peptide. Various modes of interactions between the peptide and Ca^2+^ ions were observed in these simulations by monitoring: i) distances between the Ca^2+^ ions in solution and the side chains of all acidic residues in the peptide; and ii) pairwise distances between the side chains of all acidic residues (stable binding of Ca^2+^ is expected to bring the side chains of the coordinating residues close to each other).

As shown in Figure 2A, we observed simultaneous association of 2 Ca^2+^ ions with different pairs of residues in the FP (see red and blue rectangles) in 4/18 trajectories (Replica IDs 6, 9, 14, and 16). In 3/18 simulations (Replica IDs 1, 10, and 17 in Figure 2A), binding of only one Ca^2+^ ion per FP was observed, with various pairs of residues involved. In the remaining trajectories we found instances in which a Ca^2+^ ion was associated with a single acidic residue in the FP (the events marked by grey-striped rectangles in Figure 2A). Since in such cases the coordination of the bound Ca^2+^ is not optimal and leads to low affinity/stability, these systems were not considered for further analysis.

**Figure 2:**
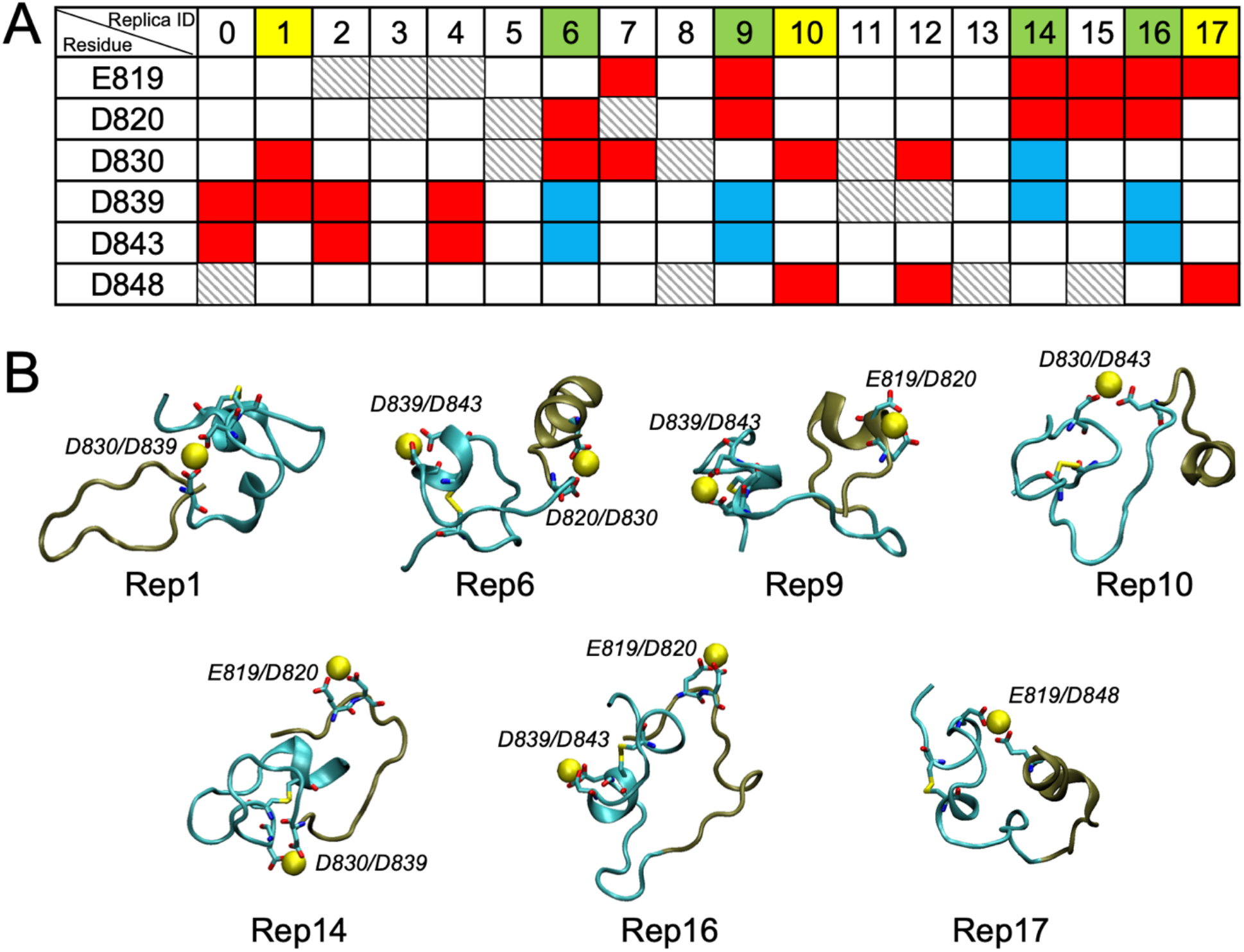
(**A**) Modes of Ca^2+^ binding to SARS-CoV-2-FP. The table shows acidic residues implicated in Ca^2+^ binding in 18 independent atomistic MD simulations (each 640ns in length). In a particular trajectory, residue pairs simultaneously engaging with the bound Ca^2+^ are denoted by red or blue rectangles, whereas instances of Ca^2+^ ion associating with a single acidic residue is depicted with grey-stripped rectangle. The trajectories in which simultaneous binding of two Ca^2+^ ions to different pairs of residues were observed are highlighted in green (Replicates 6, 9, 14, and 16). The simulations in which a single Ca^2+^ ion was bound to a pair of acidic residues are shown in yellow (Replicates 1, 10, and 17). (**B**) Structural snapshots of Ca^2+^-bound SARS-CoV-2-FP from the simulations in panel A highlighted in green and yellow (labeled according to replica ID). In the snapshots, bound Ca^2+^ ion is depicted as yellow sphere, and the N-and C-terminal parts of the fusion peptide (termed FP1 and FP2, respectively) are drawn in tan and cyan colors, correspondingly. The acidic residues engaged with Ca^2+^ ions are shown in licorice and labeled.

Overall, the binding stoichiometry in the simulations was ~1.8, in agreement with the experimentally measured value of ~1.9 for SARS-CoV-2 ^30^ and ~1.7 for SARS-CoV ^31^. Furthermore, studies of SARS-CoV have suggested a model ^31^ whereby one Ca^2+^ ion binds to the N-terminal part of the peptide (FP1, residues 801-817, corresponding to 816-835 in SARS-CoV-2), whereas another binds at the C-terminal end (FP2, residues 818-837, corresponding to 836-854 in SARS-CoV-2) – see Figure 1A. In this respect it is interesting to note that among the seven simulations of SARS-CoV-2-FP in which Ca^2+^ ions engaged with pairs of acidic residues (replicates highlighted in green and yellow in Figure 2A), the most common associations by far involved the pairs of residues E819/D820 and D839-D843, each of which occurs 3/7 times. D830 was involved in 4/7 replicas but in different pairs (D830/D839, D820/D830, and D830/D848), and D848 was observed in 2/7 replicas in different pairs (E819/D848 and D830/D848). Structural representations of these different modes of Ca^2+^ binding are illustrated by the snapshots shown in Figure 2B. From the time evolution of the secondary structure along the FP sequence in the concatenated trajectory from the replicas of FP simulations (Figure S1A in the Supplementary Material), we see several regions in which the helical structure is sustained for longer times (the N-terminus, and the 835-845 residue stretch) while folding/unfolding occurs in others. This is evident in Figure S1B showing the fraction of frames in which a particular FP residue is part of a helix. We note that these findings agree well with recently published results from CD experiments on SARS-CoV-2-FP showing that the FP is mostly unfolded in solution ^30^.

### Association of the Ca^2+^-bound SARS-CoV-2-FP with the lipid membrane

We investigated with MD simulations the spontaneous binding to lipid bilayers of each of the seven structural models of the Ca^2+^-bound SARS-CoV-2-FP in Figure 2B. Each of these structures was placed initially next to a membrane bilayer model composed of 600 lipids in a 3:1:1 mixture of POPC/POPG/Cholesterol that mimics the lipid composition used in recent experimental Ca^2+^ binding assays in cognate systems ^30,34^. For each of these seven starting points the MD simulations were run in 6 independent replicates of 1 μs length, amounting to 42μs total run time (see Table 1). As described in Methods, a set of separate control MD simulations of the SARS-CoV-2-FP with no Ca^2+^ ions bound was run with the same lipid membrane under otherwise identical conditions.

Figure S2A shows the frequency of associations with the lipid membrane for each residue of SARS-CoV-2-FP in the 42 atomistic MD simulations of the Ca^2+^-bound peptide. This measure was calculated separately for each group of 6 MD runs initiated from the same replica (identified by its Replica IDs from Figure 2B) and illustrates the preferred modes of peptide-membrane association. A residue was considered to be in contact with the membrane if the z-directional distance between its C_α_ atom and the neighboring lipid phosphorus atoms (P-atoms) was < 4Å (see Figure S2 captions for more details).

The contact frequency results show that the modes of association with the membrane involve different structural segments in the 7 replicas that represent the different modes of Ca^2+^ binding shown in Figure 2B. Thus, Rep6 and Rep9 engage with the bilayer mostly via two segments: the middle region of the peptide (residues 830-840) and the C-terminal part (see also Figure S3). Rep10 and Rep14 show associations through the middle region (residue 825-835) and the N-terminal segment, respectively. Rep16 engages with the membrane through multiple regions (N-terminal, middle, and C-terminal). Lastly, Rep1 and Rep17 exhibited relatively weak associations with the bilayer.

Interestingly, in all cases, the peptide segments interacting with the membrane contained residues that were coordinating Ca^2+^ ions (Figure S3) suggesting that the Ca^2+^ ions enhance the interactions. In turn, MD simulations of the SARS-CoV-2-FP-membrane system without bound Ca^2+^ ions registered only sporadic and much weaker binding of SARS-CoV-2-FP to the membrane (Figure S4).

To obtain a measure of the timescales of the interactions characterized in Figure S2, we first converted all trajectories into time-series of bound and unbound states (Figure S5, panels A-G) and then calculated lifetimes for each bound state. The distribution of the lifetimes (Figure S5, panel H) identified relatively long-lived complexes, ~850ns and ~770ns, respectively in trajectories from Rep9 and Rep14 (marked by red and orange stars in Figure S5). To understand the reason for the observed long lifetimes in these trajectories, we analyzed in more detail their modes of insertion into the lipid bilayer.

### Two major modes of membrane penetration by Ca^2+^-bound SARS-CoV-2-FP

The extent of peptide penetration into the lipid bilayer was quantified using two complementary approaches. In one, we monitored z-coordinate of the Cα atom of each residue in the SARS-CoV-2-FP and considered a residue inserted into the membrane if the z-distance between its C_α_ atom and the second carbon atom in the tail of a POPC lipid (atom C22 in CHARMM36 notation) was <5Å. Using this protocol, we found that the simulations initiated from Rep9, Rep10, and Rep14 resulted in the highest frequency of insertion (Figure S2B), driven by either the middle segment of the peptide (as in Rep10 and Rep14) or the N-terminal segment (in Rep9). The simulations initiated from the other replica models, either did not show insertion (Rep1), or indicated only transient membrane penetration by the C-terminal segment of the FP (Rep6, Rep16, Rep17). The latter mode of insertion is not feasible mechanistically in the context of the SARS-CoV-2 spike protein because only the N-terminal end of the FP is freed by the S2’ cleavage while the C-terminal end of the FP remains part of the spike.

Importantly, our analysis revealed that the trajectories in which the membrane-bound state is most persistent (see Figure S5, runs marked by red and orange stars) also achieve the largest extent of membrane insertion (Figure S2C, trajectories 13 and 29), but in two distinct modes of membrane penetration seen in Figure 3A and Figure S6. In one (Mode 1), the peptide inserts the F833/I834 pair of hydrophobic residues. In the other (Mode 2), the bilayer is penetrated by a different pair of hydrophobic residues, L822/L823 in the N-terminal segment of the peptide.

**Figure 3:**
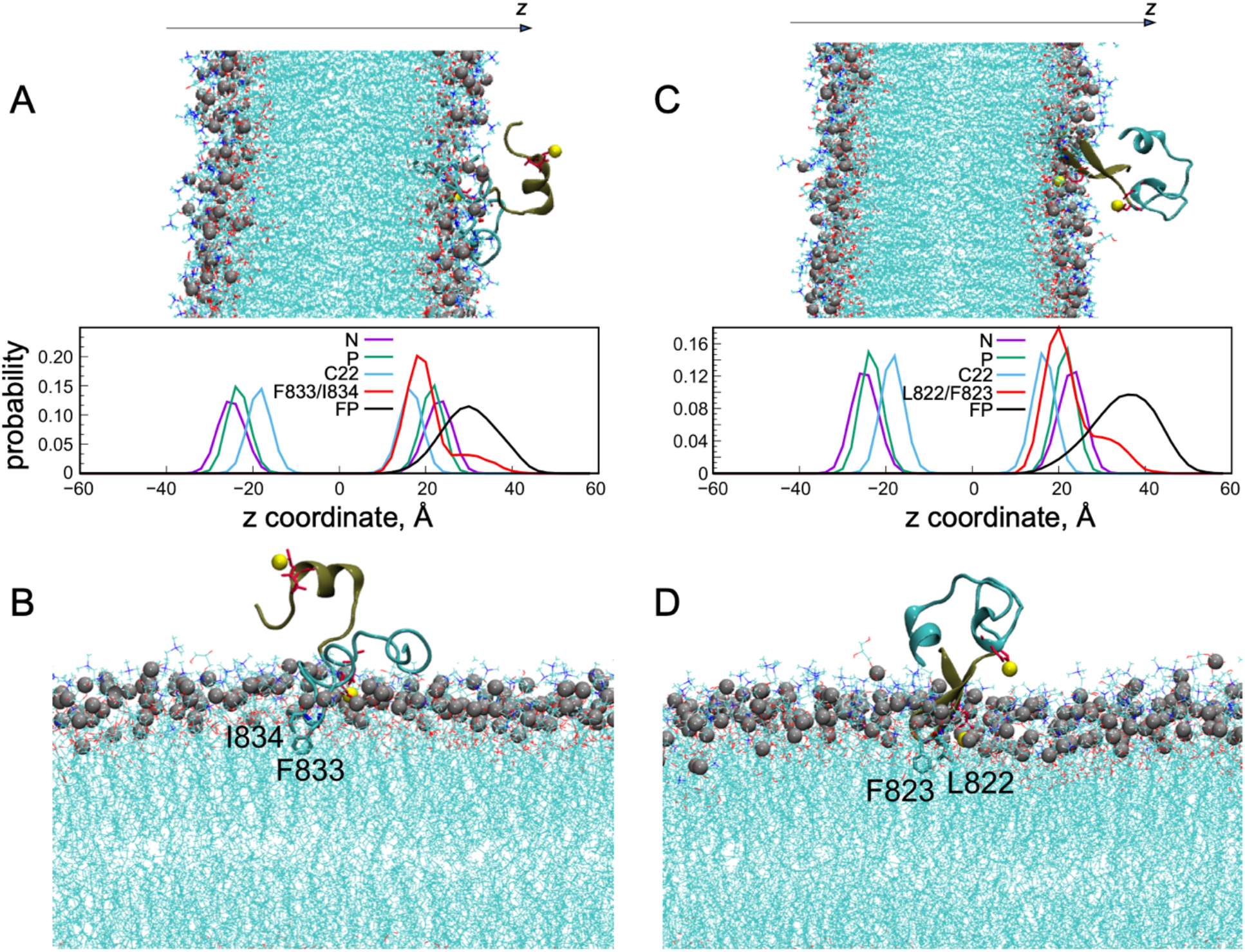
(**A**) Plots of the calculated probabilities for finding: a POPC lipid head-group nitrogen atom (N), a phosphorus atom (P), a lipid tail C22 carbon (C22), any atom of the SARS-CoV-2-FP (FP), and of the F833/I834 pair, inside 2Å rectangular slabs along the membrane-normal z axis. The data were generated from analysis of one of the MD trajectories where F833/I834 residue pair inserted into the lipid bilayer. The snapshot shows the peptide-bound bilayer in the orientation that matches the direction of the x-axis of the plots in the panel below. The P-atoms are shown as silver spheres, and the rest of the non-hydrogen lipid atoms as lines. (**B**) Close-up view of the system from panel A illustrating insertion of F833/I834 residue pair (in licorice and labeled). Color-code of the peptide is the same as in Figure 2B. The acidic residues engaged with Ca^2+^ ions (yellow spheres) are rendered in red sticks. (**C**) Plots of the calculated probabilities listed in (**A**), but for the L822/F823 pair. The data for the plots were generated as in (**A**). (**D**) Close-up view of the system from panel **C** illustrating insertion of L822/L823 residue pair (in licorice and labeled). Color-code is the same as in panel **B**. See also Figure S3 in the Supplementary Material.

To establish the specific depth of these insertions in relation to lipid head group and backbone regions, we calculated the number densities of all atoms of the peptide along the membrane-normal z-axis (i.e., the probability of finding the peptide atoms within rectangular slabs along the z-direction, see description in Figure S7 legend). These probability distributions were then compared to those for atoms of the POPC lipids at different positions, e.g., the headgroup nitrogen and phosphorus atoms (N-and P-atoms, respectively), the lipid tail second carbon atoms (C22-atoms) to identify the depth of penetration relative to the membrane thickness. Figure S7 shows that in many trajectories the density of protein atoms overlaps with that of the lipid head group atoms, consistent with our results in Figure S2A that the peptide adsorbs on the lipid bilayer. But, in addition, the analysis shows that in the trajectories of replicas exhibiting deep penetrations and the longest lifetimes of bound states, the protein density overlaps with that of the C22-atoms (trajectories 13 and 29, marked by red and yellow stars in Figure S7). This indicates that the peptide penetrates relatively deeper into the hydrophobic region of the bilayer.

Investigation of the z-directional density profiles of each residue of the peptide confirmed the deep insertion of the same residues identified from the insertion count analysis described above, namely, F833/I834 in trajectory 13 and L822/F823 in trajectory 29. This is shown by the overlap of densities of the F833/I834 pair (in Figure 3A for trajectory 13) and the L822/F823 pair (in Figure 3C for trajectory 29) with those of the lipid N-, P-, and C22-atoms. Thus, we find that the inserted hydrophobic residues partition between the P and C22 atoms of the membrane, approximately equidistantly (i.e., ~3-4Å from each of them) along the membrane normal z-axis. That these pairs of SARS-CoV-2-FP residues partition inside the membrane on the level of the C22-atoms (see also the structural snapshots in Figures 3B and 3D), agrees with results from electron spin resonance (ESR) measurements of membrane interactions of FPs from SARS-CoV ^31^, MERS-CoV ^32^, Ebola virus ^34^, as well as from the recent findings for SARS-CoV-2 ^30^. The experimental results showed that the lipid bilayer organization is perturbed in the area of the lipid head groups and backbone, but that the perturbation does not spread to the middle of the hydrophobic core of the membrane (see below for more details). That the major mode of membrane insertion for the SARS-CoV-2-FP involves the pair of residues L822/F823 is of particular importance since these belong to the highly conserved LLF motif that was shown to be critical for interactions with lipid membranes and for infectivity of SARS-CoV-FP ^31^.

Interestingly, we found that the L822/F823 membrane insertion of the Ca^2+^-bound FP was accompanied by a structural change in the N-terminal region which adopts a beta sheet conformation (Figure S8). This finding agrees with the findings from the recently published results of CD experiments showing a (4-7%) probability of the N-terminus of the SARS-CoV-2-FP folding into an extended conformation in the presence of lipid vesicles ^30^.

### Modes of membrane penetration relate to Ca^2+^ binding positions in the SARS-CoV-2-FP

The observation that the two main insertion modes were established from trajectories starting with the SARS-CoV-2-FP stabilized by the binding of two Ca^2+^ ions – one to the N-terminal residue pair E819/D820, and the other to residue pairs in the C-terminal part D839/D843 (Rep9), or D830/D839 (Rep14) – prompted a closer look at the relationship between the Ca^2+^ binding poses in the FPs and the modes of membrane insertion. Since the number of insertion events observed was rather small (Figure S6), we constructed an enhanced set of simulations to probe the interaction of *multiple* SARS-CoV-2-FPs with the lipid membrane. Simulations of such constructs consider the locally elevated concentration of the fusion peptides when one or more trimeric spike protein associate with the host cell membrane and have the advantage of enhancing the conformational sampling of the system.

The size of the constructs for these simulations was guided by the experimental conditions in studies in which the cognate SARS-CoV-2-FPs ^30^ and SARS-CoV-FPs interactions with lipid membranes were investigated for a range of peptide concentrations ^31^. Constructs were therefore built with 3, 6, 9, and 12 SARS-CoV-2-FPs (the highest concentration used in the experiments, a 2% peptide-to-lipid molar ratio ^30,31^ corresponds to the 12 FPs construct). To build these constructs, replicas of Ca^2+^-bound SARS-CoV-2-FP structures were selected randomly from the seven models shown in Figure 2B and placed on the same lipid membrane that was used for the single SARS-CoV-2-FP simulations at random positions of a rectangular grid. The starting configuration for each peptide concentration consisted of 10 different starting grids (Sim0 to Sim9) populated as detailed in Figure S9B for the 12 peptide construct structures illustrated in Figure S9A. Each of these starting conditions was simulated in 6 replicates as detailed in Table 1 and Methods.

The simulations of multiple (3, 6, 9, and 12) SARS-CoV-2-FPs interacting with the membrane revealed instances of at most 4 FPs simultaneously inserting into the lipid bilayer, but most frequently observed were concurrent insertions of 1-2 FPs (Figure S10). Importantly, we found that the mode of membrane penetration by SARS-CoV-2-FP via its N-terminal segment (Mode 2 insertion described above) was the most prevalent in these sets of simulations. This is illustrated in Figure S11 which presents the fraction of trajectory time in which each residue of the peptide is inserted into the membrane in the combined trajectories from the 3, 6, 9, or 12 FP constructs. As shown in Figure S11, these simulations captured mostly the same insertion modes observed for the single SARS-CoV-2-FP construct (Mode 1, Mode 2 and the C-terminus segment insertion which is possible only in the truncated FP construct). The highest frequency of insertions in all systems was found to involve the L822/F823 pair of residues (i.e., Mode 2).

To examine the relationship between the observed modes of membrane insertion and the initial structural models in the large number of trajectories from the multiple-PF constructs with randomly selected initial structure, we first quantified membrane insertions separately for those peptides that shared the same initial structure (irrespective of their initial placement on the grid). The histograms of membrane insertions for the 7 separate sets of data corresponding to the seven initial models are shown in Figure 4A. The striking observation from this comparison is that Mode 2 insertion for Rep14 model appears in all 4 sets of simulations (3, 6, 9, and 12 FPs) and results exclusively from Rep14 and Rep1 models, regardless of peptide concentration (see also snapshots in Figure 4B-C). Notably, out of the seven models in Figure 2, only Rep14 and Rep1 include a Ca^2+^ ion bound to the D830/D839 pair of residues in the FP2 segment of the peptide. pair). Together, these results suggest that the insertion of the N-terminal segment and the conserved hydrophobic motif residues is associated with a specific pattern of Ca^2+^ ion binding to the SARS-CoV-2 FP. This specific pattern involves the E819/D820 and D830/D839 pairs of residues which facilitate the prevalent Mode 2 of SARS-CoV-2-FP insertion.

**Figure 4:**
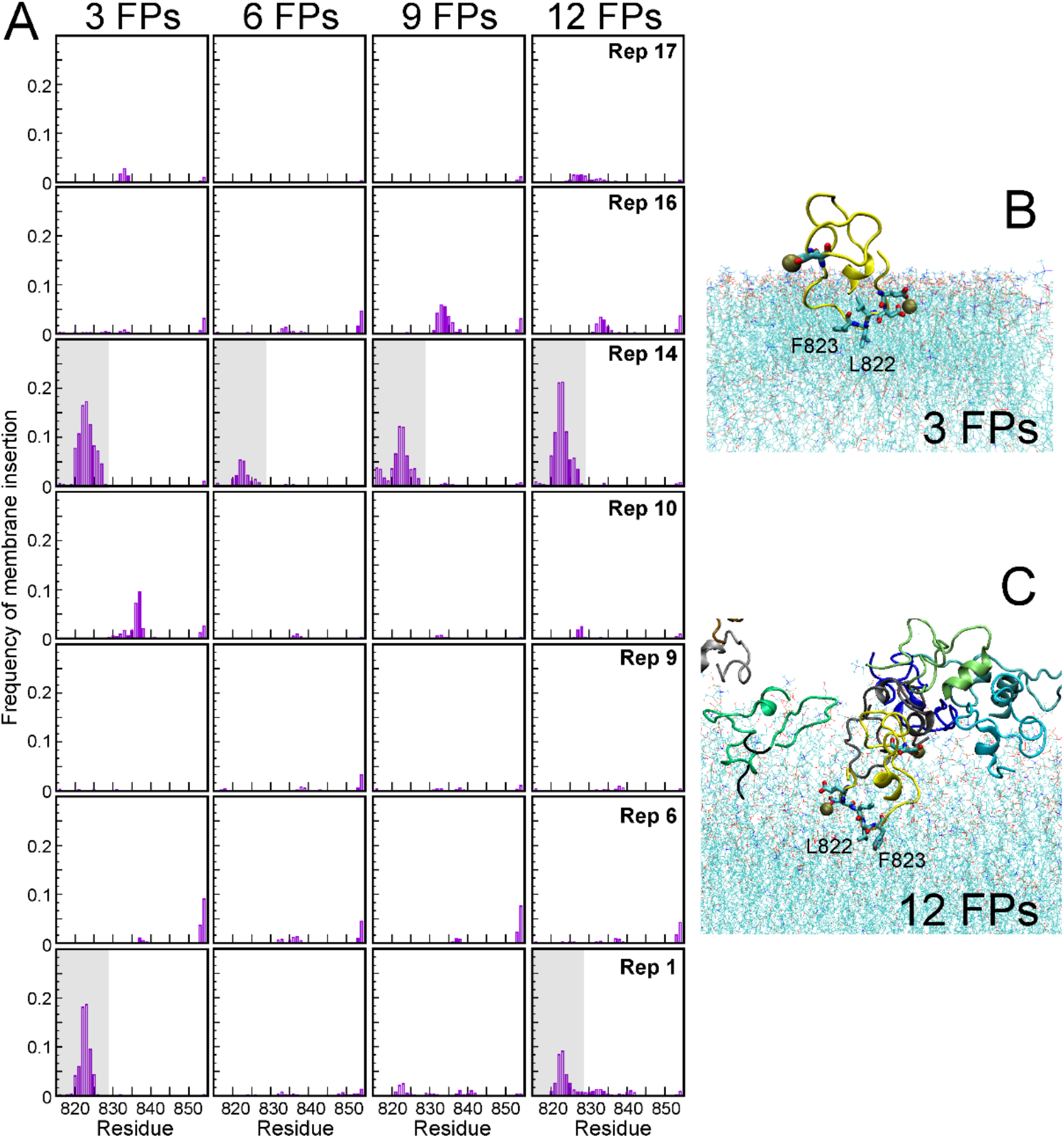
(**A**) The frequency of membrane insertion calculated for each residue of SARS-CoV-2-FP from the MD simulations of 3, 6, 9, and 12 FPs interacting with 3:1:1 POPC/POPG/Cholesterol bilayer. The data shown in each row is an average over each of the 6 starting models (Rep-s) of the peptide, identified as in Figure 2B. Membrane insertion was defined as described in the caption of Figure S2B. The conditions that resulted in the insertion of the N-terminal segment of the peptide are highlighted with colored boxes (Rep 1; Rep14). (**B**&**C**) Representative snapshots of FP insertion in the 3 FP and 12 FP systems (panels B and C, respectively) illustrating the insertion of the N-terminus segment into the membrane. The FP which penetrates the bilayer is in yellow. The Ca^2+^ ions bound to this FP are shown as golden spheres. The residues coordinating the Ca^2+^ ion are rendered in licorice. The L822/L823 pair which inserts into the bilayer is rendered in licorice and is labeled.

### Response of the membrane environment to the prevalent mode of SARS-CoV-2-FP insertion

To probe the arrangement of the individual membrane constituents around the peptide, we tracked the number of POPC, POPG, and cholesterol lipids in the bilayer near the FP in the single peptide construct trajectories where Mode 2 of insertion was stable (see Figure 3C-D). As shown in Figure 5A, the environment within 4Å of the peptide is enriched in POPG and cholesterol, specifically around the N-terminal 820-827 segment, so that the compositions of these two components locally near the peptide increases from the average 0.2 to ~0.3, while the composition of POPC decreases from 0.6 to ~0.4 (Figure 5B). A stabilizing effect of these changes is suggested by the appearance of an electrostatic association of the negatively charged POPG lipid headgroups with the basic sidechain of residue K825 which is juxtaposed to the 819/820 Ca^2+^ binding site (Figure 5C).

**Figure 5:**
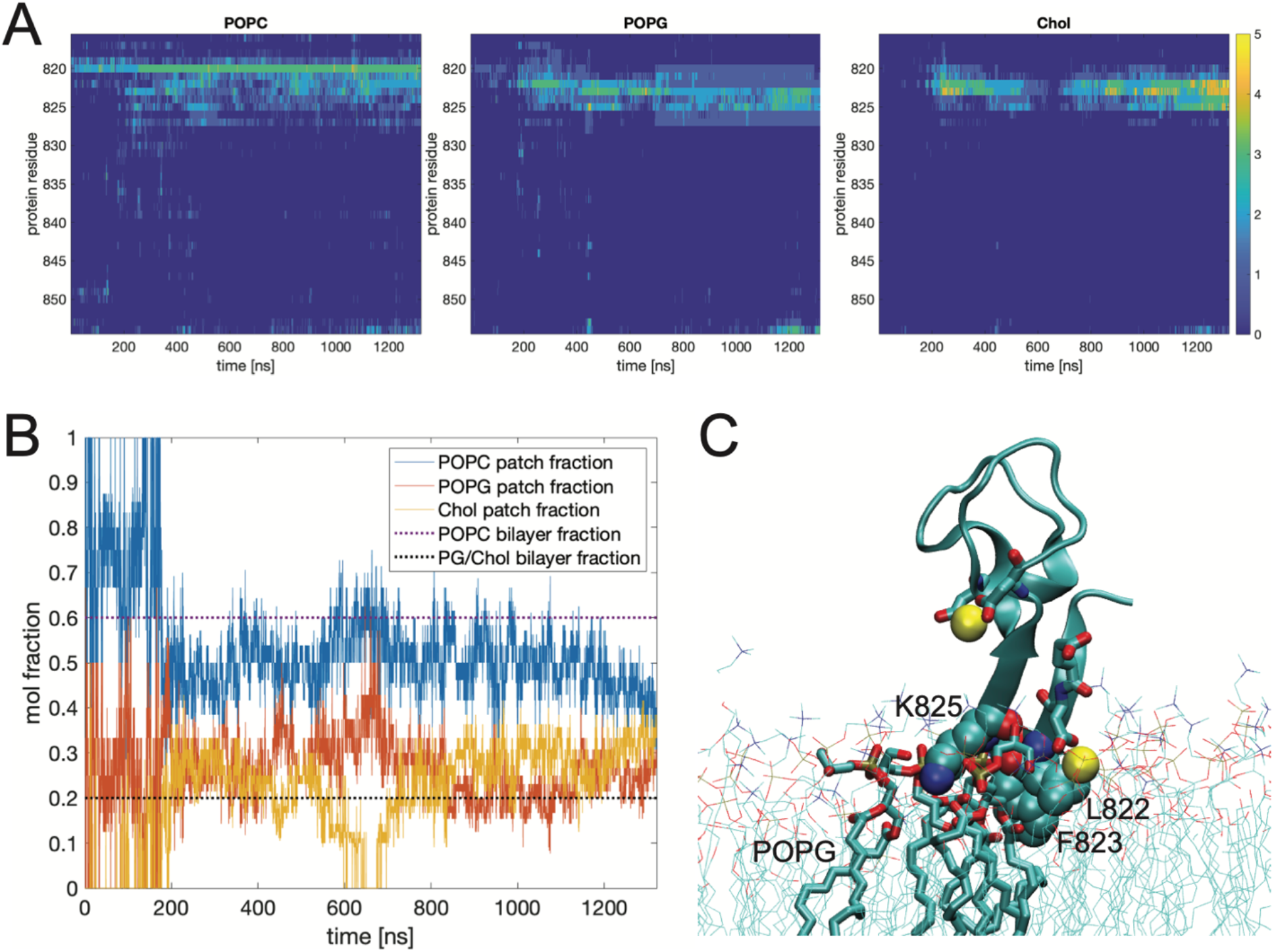
(**A**) Time-evolution (y-axis) of the number of contacts (i.e., within 4Å) per SARS-CoV-2-FP residue (x-axis) with POPC (left), POPG (middle), and cholesterol (right) in the simulations of the single FP inserted into the membrane via the LLF motif (see also Figure 3C-D). (**B**) Time-evolution of the fraction of POPC (blue), POPG (red), and cholesterol (yellow) molecules in contact with the peptide in the same MD trajectory as in panel A. A lipid was considered in contact with the FP if the 3D distance between any lipid atom and any FP atom was < 4Å. Dotted lines indicate the bulk compositions of these lipids in the mixture (3:1:1 POPC/POPG/Cholesterol). (**C**) A snapshot of the system (as in Figure 3D) highlighting POPG lipids (in licorice) within 4Å of the peptide. Residues L822, F823, and K825 are shown in space fill and are labeled. The two bound Ca^2+^ ions are shown as yellow spheres, and the residues coordinating them are drawn in licorice.

The effects of these rearrangements in local composition, and the presence of Ca^2+^ ions being able to interact with the membrane and the hydrophobic residues at the different levels of the proximal bilayer leaflet, are expected to modify membrane properties locally and be propagated more distantly. To evaluate the consequences of the observed increase in the local cholesterol concentration around the peptide we calculated the ordering effect on the lipids in the vicinity of the insertion in the radial zones defined in Figure 6A. The calculated deuterium (S_CD_) order parameters for POPC and POPG lipid tails as well as orientations of PC and PG headgroups in the different zones are shown in panels B-E of Figure 6. For the lipids situated in the zones closest to the insertion we find higher SCD values indicating higher order in their hydrocarbon chains that propagates to ~30Å from the insertion (Zones 1 and 2). Interestingly, we also find the PG headgroups (but not PC headgroups) in more vertical, “upright” orientation near the peptide (indicated by lower angle values with respect to membrane normal shown in panels 8F and 8G). This headgroup ordering effect is likely due to the observed electrostatic interactions between the PG head groups and K825 as it is confined to the first zone closest to the FP.

**Figure 6:**
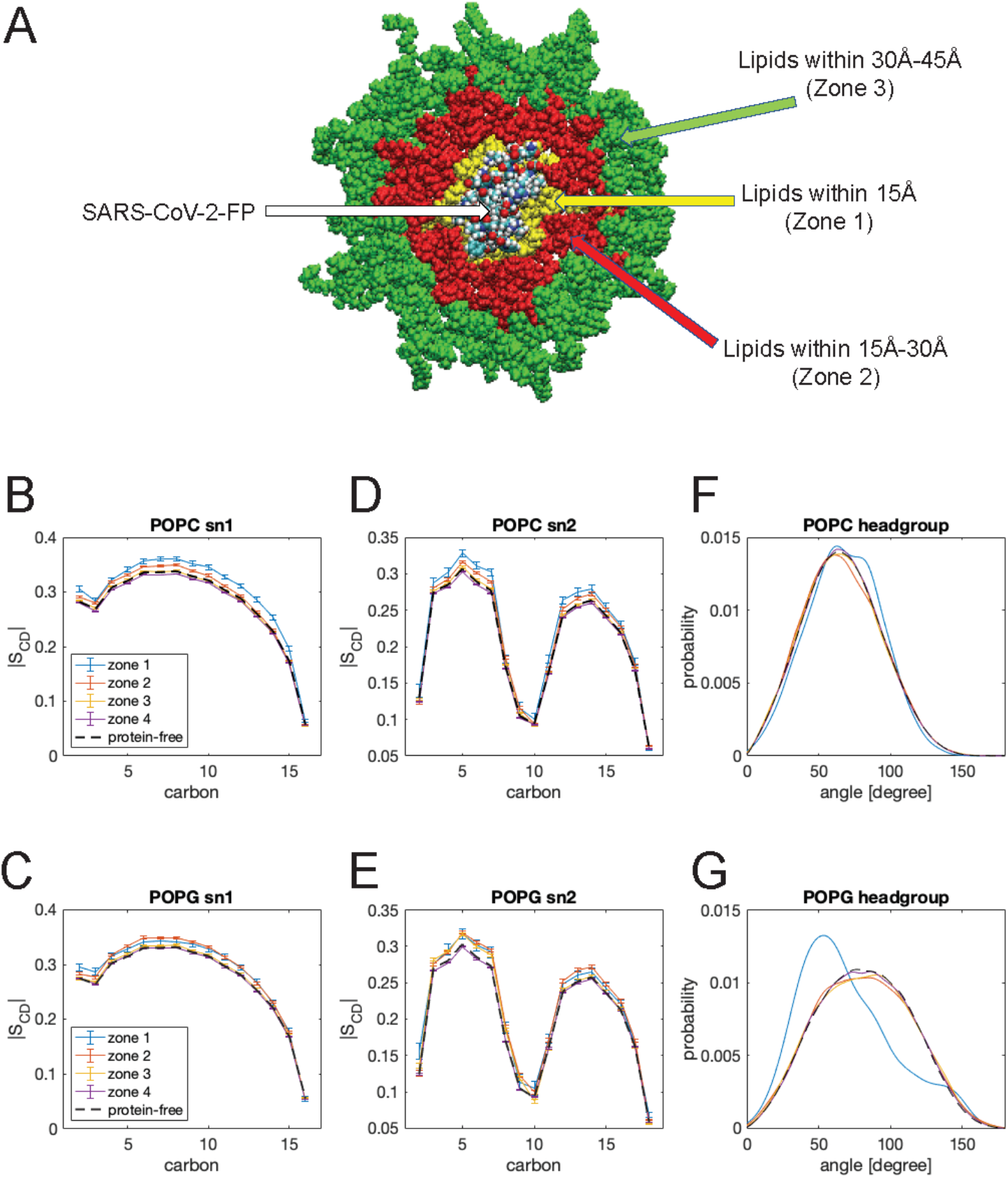
(**A**) Definition of radial zones in the membrane around the inserted SARS-CoV-2-FP. A lipid was considered to be in Zone 1, 2, or 3 if its phosphorus atom was within the lateral distance of 15Å, 15Å-30Å, or 30Å-45Å, respectively, from the center of mass of the FP. All other lipids constitute Zone 4. (**B**-**E**) Deuterium order parameters for carbon atoms in the sn1 (**B**, **C**) and sn2 (**D**, **E**) hydrocarbon chains of POPC (**B**, **D**) and POPG (**C**,**E**) lipids calculated in the zones defined in **A**. The analysis was carried out on the trajectory of 1FP-membrane system (+1FP) that was used also for the analysis of the pressure profile (see text, also Figure 7). For reference, order parameters of the protein-free system are shown in dashed lines. (**F**-**G**) Orientation of the POPC (**F**) and POPG (**G**) lipid headgroups (P-N, and P-C12 vectors, respectively) with respect to the bilayer normal calculated in the different zones defined in panel **A**. The analysis was carried out on the same trajectory used in **B-E** and for pressure profile calculations Figure 7).

We next investigated if these FP-induced local structural perturbations in the membrane induce more global effects on bilayer properties, such as on the lateral pressure distribution ^45^. Pressure profiles were calculated (see Methods) from the single FP trajectories of Mode 2 insertion (Figure 3C-D), and from one of the replicates from 12 FPs construct in which 2 FPs were stably inserted via Mode 2 (Figure S12). The calculated lateral pressure profiles as a function of the membrane-normal z coordinate in these two trajectories are shown in panels Figure 7A&B (red lines). For comparison, the black lines in these panels show the pressure distribution calculated for the peptide-free membrane system. To aid the interpretation of these plots in the structural context of the bilayer and the peptide, Figure 7C&D shows the position densities along the membrane normal for selected lipid atoms (P, N, and C) and FP atoms.

**Figure 7:**
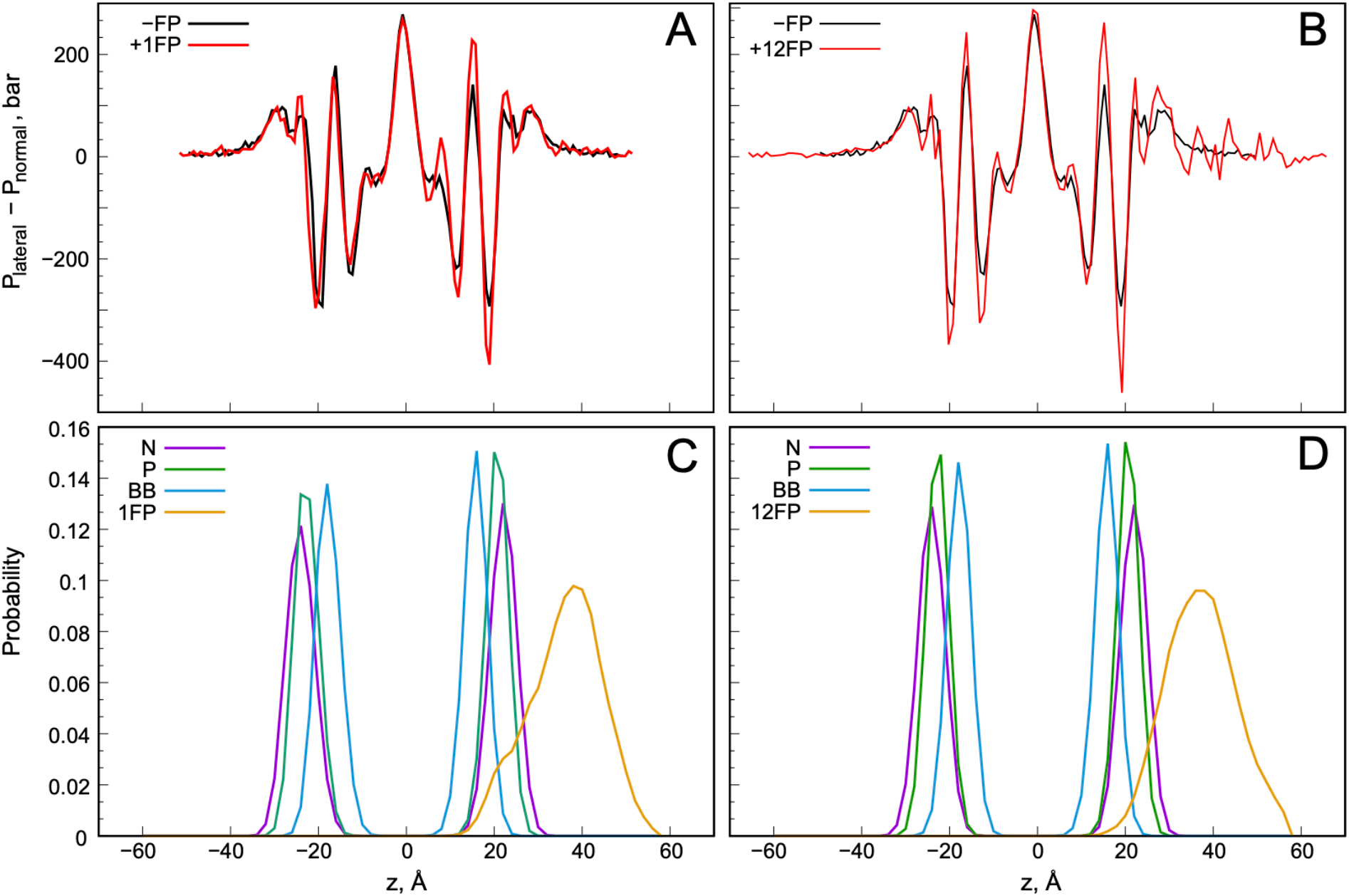
(**A**-**B**) The pressure profile as a function of the membrane-normal z coordinate in the simulations of a single FP (+1FP, panel **A**) and 12 FPs (+12FP, panel **B**). For comparison, the pressure distribution in the peptide-free system is also shown (-FP). (**C**-**D**) Plots of the probability of finding POPC lipid head-group nitrogen atom (N), phosphorus atom (P), hydrocarbon chain C22 carbon (C22), and any atom of the SARS-CoV-2-FPs (FP) in 2Å rectangular slabs along the membrane-normal z axis calculated from the trajectories from simulations of a single FP (1FP, panel **C**) and 12 FPs (12FP, panel **D**).

The pressure profile in the pure bilayer system follows the expected trends ^46^: in the bulk solution, where the system is stress-free, the lateral pressure is near zero. Upon approaching the membrane, the lateral pressure deviates from zero and develops both positive and negative peaks in the lipid headgroup and backbone regions (10Å<|z|<30Å in Figure 7). The negative peaks are reflective of attractive forces at the membrane/water interface that shield the hydrophobic core of the membrane from solvent exposure and thus reduce the bilayer area, whereas the positive peaks are due to repulsive forces that increase the bilayer area. The additional positive peak in the bilayer center (z=0 in Figure 7) stems from repulsive forces that arise from entropic losses in dynamics of the lipid tails ^46^.

The insertion of a single FP peptide has significant effects on the pressure distribution in the leaflet proximal to the peptide, and the profile becomes asymmetric between the two leaflets. Indeed, as shown in Figure 7A, both the positive and negative pressure peaks in the FP-inserted leaflet (z>0, see also Figure 7C) are higher than in the pure bilayer, but the pressure is unaltered at the bilayer midplane and in the opposite leaflet. These effects are consistent with the changes in the orientation of the phospholipid headgroups as well as the trends in the acyl chain order parameter profiles of POPC and POPG (Figure 6) whereby the ordering effect caused by FP is visible across most of the length of the lipid chains except for the last few carbons situated at the bilayer midplane.

The analysis of pressure profiles in the 12FP system with two FPs simultaneously inserted in the membrane (Figure 7B) reveals even more striking differences from the bare membrane system. All peptides, including the two FPs that interact with the membrane, are in the z>0 region of the simulation box, which makes the pressure in the bulk solution (z>30Å in Figure 7B) deviate strongly from zero. Importantly, while the pressure profile retains similar asymmetry as in the 1FP system, with the top leaflet exhibiting larger positive and negative peaks than the bottom one, the two inserted peptides affect the pressure distribution in the entire bilayer.

To understand the difference between the membrane perturbations in the single FP compared to the 12 FP construct, we examined the structural properties of the two leaflets separately in these two systems and compared them to those in the peptide-free bilayer. We found that the average order parameter values for POPC were similar in the two leaflets of the 12 FP construct, but lower than those in the bulk (pure) membrane system (Figure S13). This change is accompanied by a slight increase in the average area per lipid (Figure S14) and shows an overall deviation from bulk properties, indicative of the presence of stress in the membrane.

As noted earlier and shown in Figure 6, the structural perturbations induced by a single FP can propagate up to a distance of ~30Å. At the highest protein-to-lipid ratio used in experiments and simulated in the 12FP system, the distance between any two peptides bound to the membrane surface is less than 60Å (1/2 of the length in x and y directions of the simulated membrane). Thus, in the presence of multiple bound peptides the perturbation ranges overlap so that perturbed bilayer properties cannot relax to their unperturbed values. This results in the accumulation of stress across the entire bilayer. While the structural changes detected in the 12FP simulation (Figures S13–S14) are small overall, and consistent with a low bilayer tension (between 1.5 and 2 mN/m), they are expected to increase with the number of peptides binding to the membrane. Taken together, these findings show that interactions of FPs from SARS-CoV-2 (or SARS-CoV, which shares the highly conserved FP sequence) induce measurable perturbations of lipid bilayer organization and properties that are dependent on the local peptide concentration and can affect the areas of the lipid head groups, backbone, and most of the length of the lipid chains.

## DISCUSSION

In the exploration of targets for small molecule inhibitors that can prevent coronaviruses like SARS-CoV-2 from entering human cells, efforts had been directed mostly towards disrupting the molecular interaction of the receptor binding domain (RBD) of the Spike protein with the ACE2 receptor (see Refs. ^5,47,48^ and citations therein). But with the growing experimental evidence for the involvement of the FP region of the spike protein in the interaction with the cell membrane during the entrance of the virus ^22^, and role of this region in the fusion of the cell and viral membranes ^5^, we considered the FPs as another possible target for efforts to identify and/or design effective therapeutics against CoVID-19 disease. However, unlike the RBD/ACE2 interactions of the SARS and MERS corona viruses, which have been mapped out from structural data at the atomistic level, the structure-function relations underlying the molecular mechanisms of their FP regions in the fusion-ready state of the Spike protein, lacks this level of definition and understanding. The structure of the fusion peptide in the context of the Spike trimer of SARS viruses has been determined to some extent. Indeed, the highest resolution structure of the SARS-CoV-2 Spike in this state ^23^, offers interesting clues for a role of the FP2 region (referred to as the “fusion peptide proximal region”, FPPR) in preparing the recognition of the RBD by the ACE2 receptor. But there is no information about the structure or dynamic properties of the FP in the penetration and “post fusion” state ^23^ of the Spike. The computational analyses presented in this work have addressed this gap in the information needed to target the FP region of the SARS-CoV-2 virus for inhibition of membrane penetration.

Using the described analyses of the trajectories from swarms of MD simulations of the SARS-CoV-2-FP we predicted various feasible Ca^2+^-binding positions (“poses”) (Figure 2). One of the most prevalent poses that emerged from the analysis (see Figure 2B) binds two Ca^2+^ ions, one in the N-terminal region (FP1) at the E819/D820 pair, and the other in the C-terminal region (FP2), at D830/D839. Remarkably, simulations probing the spontaneous binding of the SARS-CoV-2 fusion peptide to lipid membranes (started from all the identified Ca^2+^-binding poses under conditions representing various concentrations of the FP), revealed that binding and penetration are achieved in only two major modes. Moreover, the predominant mode of interaction with the membrane involves the Ca^2+^-binding pose we had identified to be one of the most likely ones, with the ion bound at E819/D820 and D830/D839. The FP models with this Ca^2+^-binding pose insert the N-terminal hydrophobic residues L822 and F823 – which are part of the conserved LLF motif shown to be of functional importance in SARS-CoV-FP ^31^ – into the hydrophobic portion of the membrane. This predominant mode is stabilized by several factors including the electrostatic interactions of (i)-the Ca^2+^ bound to E819/D820 with the lipid headgroups, and (ii)-the anionic POPG lipid headgroups with the basic sidechain of K825 juxtaposed to the LLF segment. This stabilization suggests an answer to the question raised by the experimental finding that Ca^2+^ can enhance the interactions between the peptide and the lipid membrane and stabilize the insertion. While the participation of the charged lipid headgroups in this stabilization is interesting, more computational studies involving other anionic lipids will probe further these mechanistic inferences. Indeed, the agreement with the experimental results in the same membranes with the published data for the FP of SARS-CoV ^31^ and MERS ^32^ is very reassuring, and as shown by the recently deposited results in bioRxiv for Sars-CoV-2 FP-membrane interactions ^30^, the agreement extends to the COVID-19 agent. Notably, experimental studies using POPC/POPG/Cholesterol or POPC/POPS/Cholesterol mixtures have produced similar results regarding the strength of the FP-membrane interactions and the extent of the FP-induced membrane ordering ^30,31^, suggesting that the charge of the lipid headgroup and not its identity, is the critical parameter here.

Given its prevalence in our simulations and in light of the established role of the LLF motif in SARS-CoV membrane binding and infectivity ^19,31,33^, we propose that the membrane-insertion mode involving the L822/F823 pair is likely to be functionally relevant in creating the fusion-competent conditions. We note that as this manuscript was being finalized, we became aware of a complementary work in bioRxiv ^49^, in which the conformations of membrane-bound *apo* models of SARS-CoV-2 (i.e., in the absence of Ca^2+^ ions) are thoroughly evaluated with MD using a coarse-grained model of the lipid membrane (HMMM). In agreement with our findings, these studies also suggested a membrane-binding mode that involves the N-terminal region of the peptide, but with poses and modes of penetration that relate to the absence of bound Ca^2+^. Consonant with the experimentally demonstrated Ca^2+^ binding requirement for efficient insertion, our findings lead to a proposed mechanism for the role of Ca^2+^ in modulating FP-membrane interactions. This mechanism describes how the preferred mode of Ca^2+^ binding to the E819/D820 pair supports the preference of the N-terminal mode of insertion which puts into play the conserved hydrophobic motif. The calculated membrane perturbation by this mode of insertion, discussed further below, supports the role of this mode of insertion in the preparation of the fusion of viral and cell membranes. Our finding that this mode of interaction is stabilized by the binding of the second Ca^2+^ to the D830/D839 pair, brings to light the interesting observation that the alternative binding of Ca^2+^ to another set of acidic residues in the C-terminus, such as D839/D843 pair, inhibits the fusion-competent mode of membrane binding. Given the high degree of sequence similarity and conservation among FPs of other coronaviruses such as SARS-CoV and MERS-CoV, we propose that these insights about structure-function relations for SARS-CoV-2, are generalizable to other corona viruses.

Analyses of the effects of the SARS-CoV-2-FP insertion on the dynamic properties of the membrane patch used in the MD simulation trajectories (with a composition mimicking the experimental conditions of cognate measurements ^34^) reveals a perturbation pattern that is consistent with results from earlier studies on FPs of SARS-CoV2, SARS-CoV, MERS-CoV, and Ebola virus ^30–32,34^. We find that FP binding modifies the molecular organization in the area of the lipid head groups and backbone atoms, as well as in most of the hydrocarbon chain layer; effects on the middle region of the bilayer hydrophobic core are minimal. Interestingly, we observed an enrichment of negatively charged PG lipids and cholesterol molecules near the inserted peptide. As described above, the electrostatic association of PG headgroups with the FP appears to provide additional stability for the LLF insertion mode, while segregation of cholesterol has an ordering effect on the lipids around FP that extends to ~30Å from the insertion site (Figure 6). Given that the viral particle will bring multiple FPs to operate in close proximity to each other near the membrane, simultaneous interactions/penetration by multiple FPs will destabilize the bilayer as indicated by our observations for the 12FP system (Figure 7). Such bilayer perturbations may be essential for the fusion of the viral and host cell membranes. Future mechanistic studies, involving more complete molecular models of the Spike as well as lipid compositions closer to those of physiological cell membranes will be necessary to address these mechanistic hypotheses and to test functional implications of membrane perturbations.

At this stage, however, the results offer a structure-specific platform to aid the ongoing efforts to design inhibitors of virus cell entry at a new and mechanistically important target. Moreover, given the conservation of both sequence and cell-penetration mechanisms of specific CoVs known to be human pathogens, and the evidence that calcium binding is both essential and shared by FPs of such corona viruses, the known structural similarities and differences in the key regions of the mechanisms we describe should be very useful in seeking specific modes of blocking the penetration and infectivity of this larger set of CoVs.

## AUTHOR CONTRIBUTIONS

GK and HW designed the study. GK carried out molecular dynamics simulations. GK, AP and MD performed analyses of the molecular dynamics trajectories. All the authors interpreted results. GK and HW wrote the manuscript. All the authors participated in the reviewing/editing of the manuscript and approved the final draft.

## ACKNOWLEDGMENTS

The authors thank members of the Weinstein lab, and in particular Drs. Giulia Morra and Derek Shore, for helpful discussions. H.W. and G.K. gratefully acknowledge support from the 1923 Fund. M.D. was supported by NIH F32GM134704. The authors gratefully acknowledge the enabling roles of Chris Carothers-director of the Center for Computational Innovations (CCI) at the Rensselaer Polytechnic Institute (RPI), and Geralyn Miller, director of health strategy at the Microsoft AI for Good Research Lab and her team at Microsoft AI and Microsoft Azure HPC, for the kind help and support of efficient and sustained access to the AiMOS supercomputer at CCI, and the Microsoft Azure HPC resources, respectively. These resources were generously awarded through the COVID-19 High Performance Computing Consortium. The authors gratefully acknowledge as well the use, in early stages of the research, of resources of the Oak Ridge Leadership Computing Facility, which is a DOE Office of Science User Facility supported under Contract DE-AC05-00OR22725, and of in-house computational resources of the David A. Cofrin Center for Biomedical Information in the Institute for Computational Biomedicine at Weill Cornell Medical College.

## SUPPLEMENTAL FIGURES

**Figure S1:**
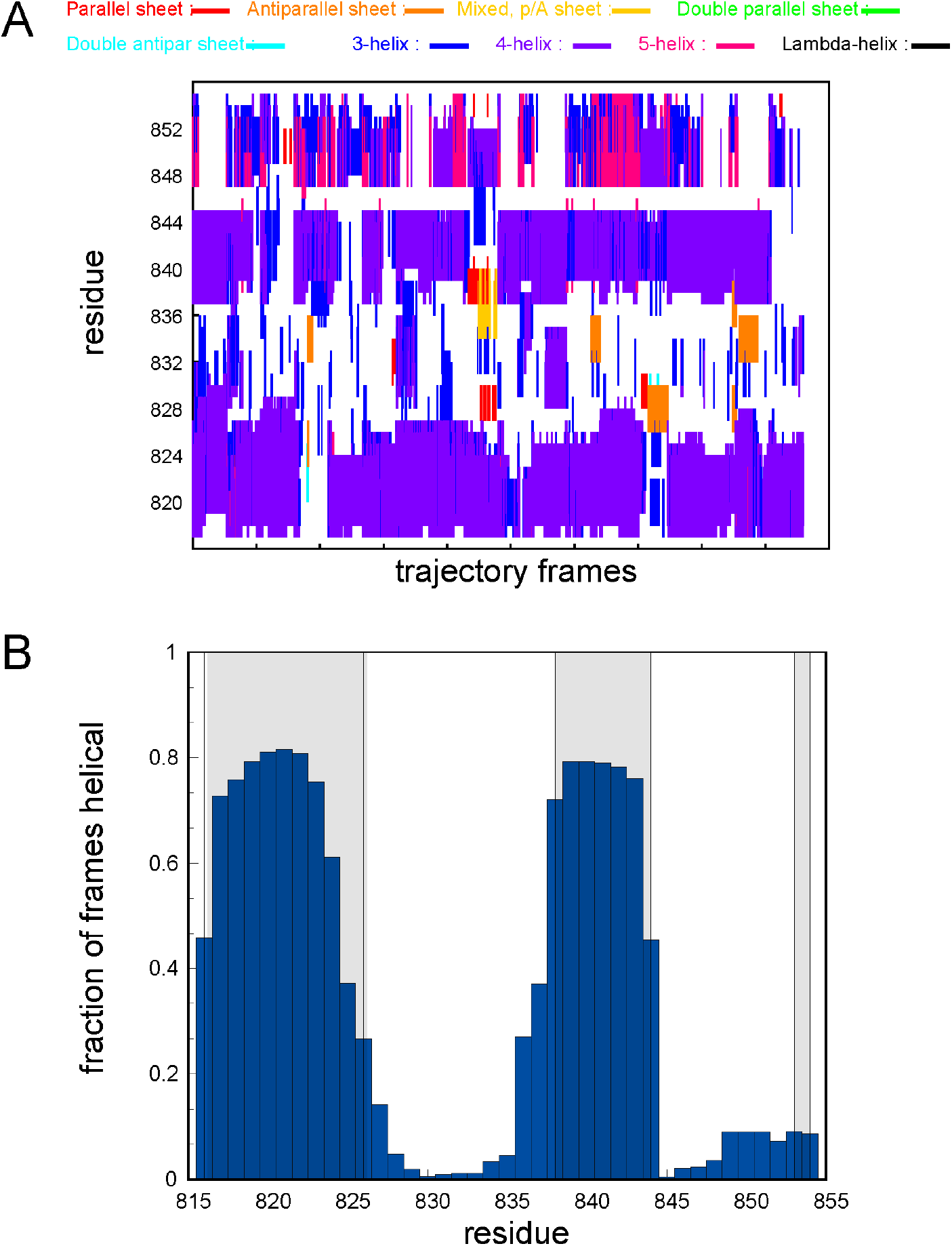
(A) Time-evolution of secondary structure elements for each Sars-CoV-2-FP residue in the simulations of the peptide in water. The analysis was performed using DSSP module in VMD on a concatenated trajectory combining the 18 independent MD replicates. (B) Fraction of trajectory frames in which a particular residue is in helical conformation. The shaded rectangles demarcate the FP regions that were helical in the initial model of the peptide.

**Figure S2:**
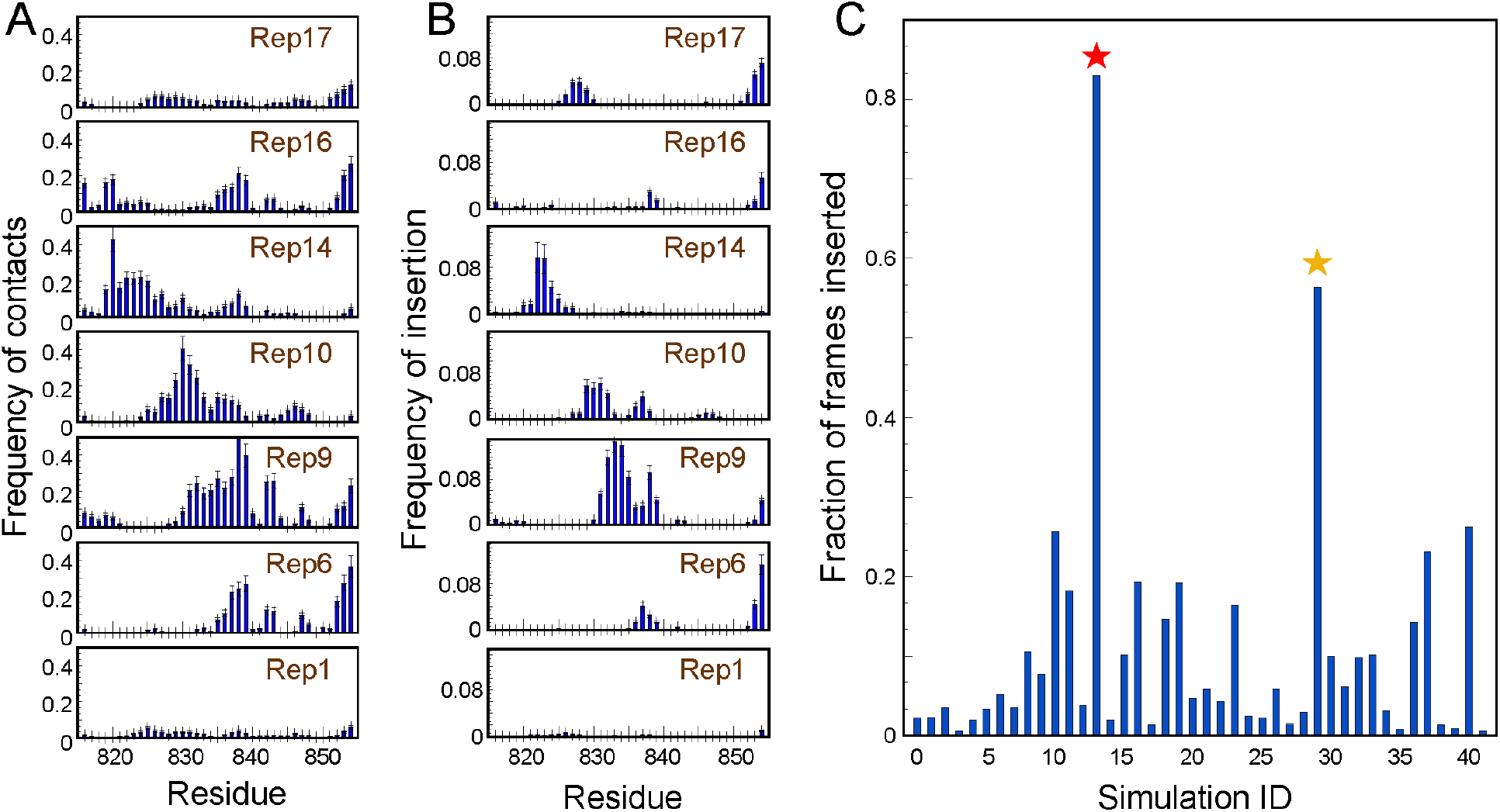
(**A**) Frequency of contacts with the lipid membrane for each residue of SARS-CoV-2-FP, calculated separately for each of the seven sets of 6 runs initiated from the common Ca^2+^-bound conformation of the peptide (see Figure 2B). Panels corresponding to different sets are labeled by the starting replica identified in Figure 2B. A residue was considered in contact with the membrane if the z-directional distance between its Cα atom and the neighboring lipid phosphorus atoms (P-atoms) was < 4Å. A P-atom was considered in the neighborhood of the Cα atom if the distance between them, projected onto the membrane x-y plane was < 10Å. (**B**) Frequency of membrane insertion for each residue of the SARS-CoV-2-FP plotted separately for the seven sets of 6 MD runs initiated from the common Ca^2+^-bound conformation of the peptide. A peptide residue was considered to be inserted into the membrane if the z distance between its Cα atom and the neighboring C22 lipid tail carbons (C22-atoms) was <5Å. A C22-atom was considered in the neighborhood of the Cα atom, if the distance between the two atoms projected onto the membrane x-y plane was < 10Å. (**C**) Fraction of frames in each individual trajectory in which the peptide was inserted into the bilayer. The peptide was considered inserted into the membrane if the z distance between any Cα atom on the peptide and the C22-atoms was <5Å. The red and yellow stars indicate Rep13 and Rep29 which are discussed in detail in the text and in Figure 4. The error bars in panels **A** and **B** were estimated from bootstrapping analysis where calculations were carried out on 20 sets of randomly chosen 6 replicates. The uncertainties were then determined from standard deviations obtained from these 20 runs.

**Figure S3:**
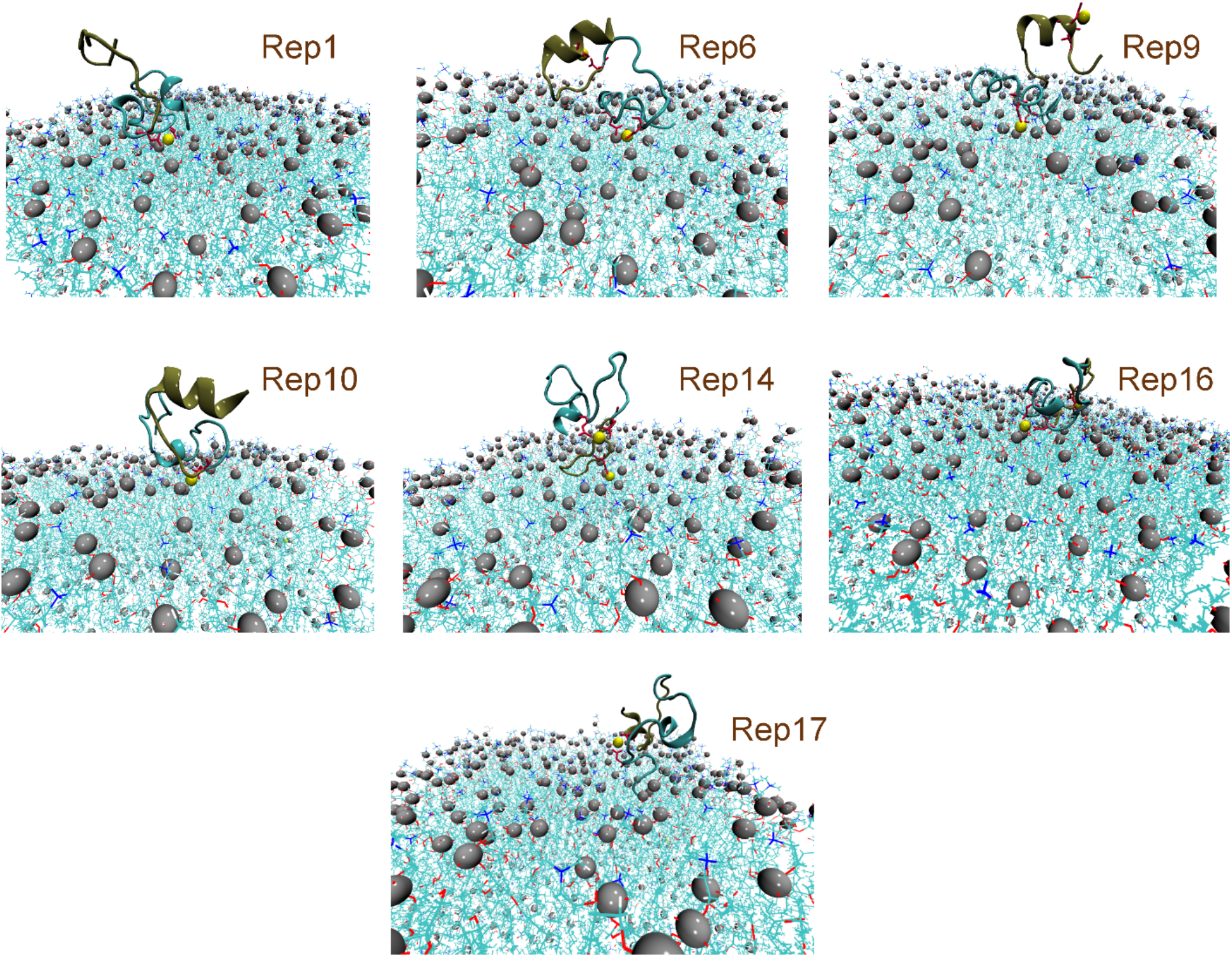
Representative snapshots from MD simulations of the Ca^2+^-bound SARS-CoV-2-FP replicas interacting with lipid membranes. The structural models shown are from each set of simulations that started from the same initial structure of the peptide and are labeled according to the scheme shown in Figure 2B of the main text. In the snapshots, Ca^2+^ ions are shown are yellow spheres. In the membrane patches surrounding the SARS-CoV-2-FP phosphorus atoms are shown as silver spheres, and non-hydrogen atoms of the lipids are rendered as lines. The N-terminal (FP1) and the C-terminal (FP2) parts of the peptide are identified in tan and cyan colors, respectively.

**Figure S4:**
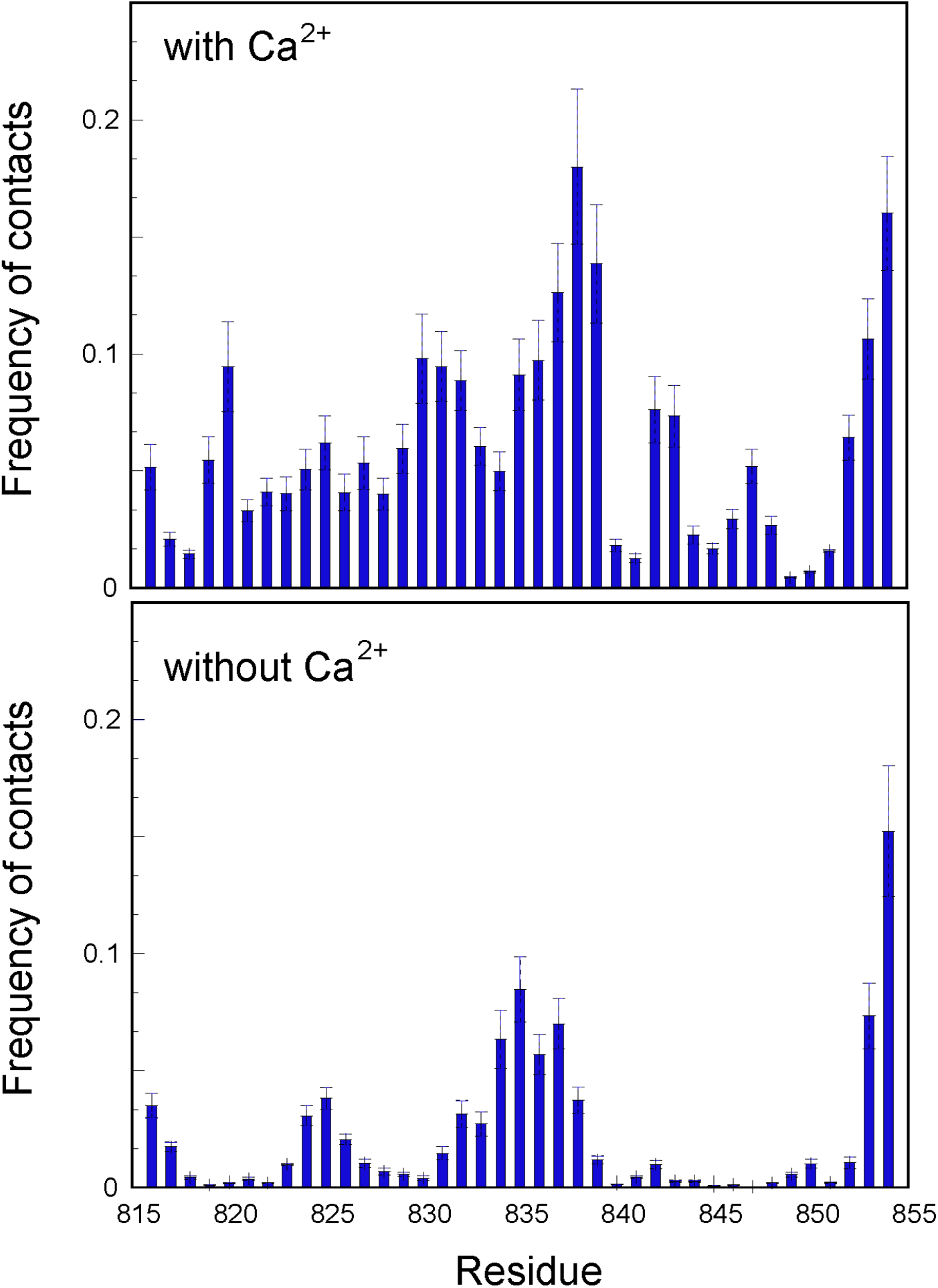
Histograms of the frequency of contacts with the lipid membrane calculated for each residue of the SARS-CoV-2-FP from the MD trajectories of the single SARS-CoV-2-FP-lipid bilayer system in the presence (*upper panel*) and absence (*lower panel)* of Ca^2+^ ions.

**Figure S5:**
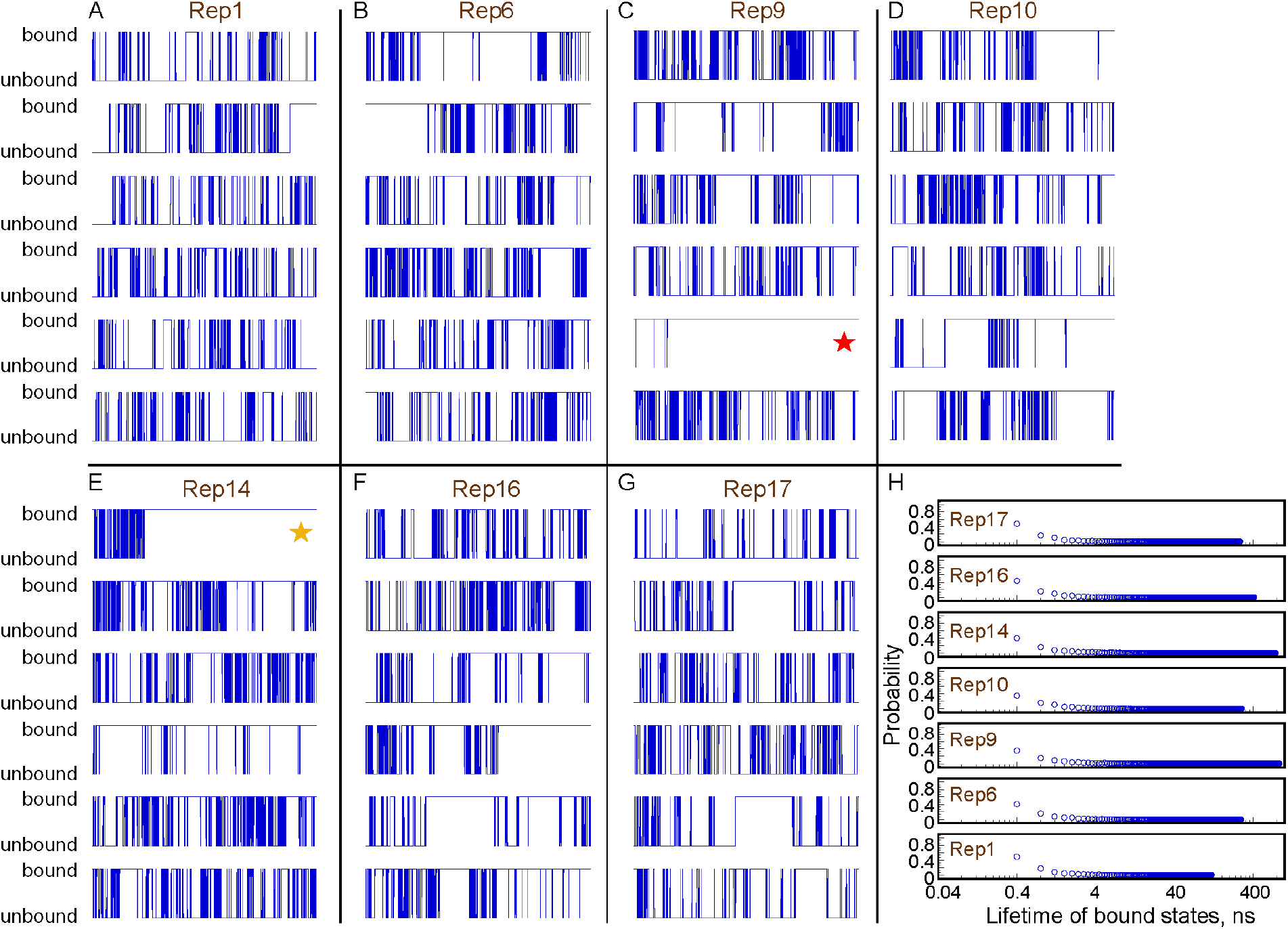
(**A-G**) Time traces from individual trajectories of the single SARS-CoV-2-FP-membrane system showing events of peptide binding to the lipid bilayer (blue lines indicate bound states). All simulations starting from the same structure of the peptide are grouped in a panel labeled according to the naming scheme shown in Figure 2B of the main text. The two trajectories with the longest lifetimes of the bound state are marked by red and orange stars and are described in detail in Figure 4 of the main text. Membrane bound states were defined as described in Figure 3 of the main text. (**H**) Lifetimes of membrane-bound conformations extracted from the time traces in panels A-G. The sets of 6 simulations per each initial condition were analyzed jointly (as labeled from Rep1 to Rep17). Note logarithmic scale on the *x* axis of the plot.

**Figure S6:**
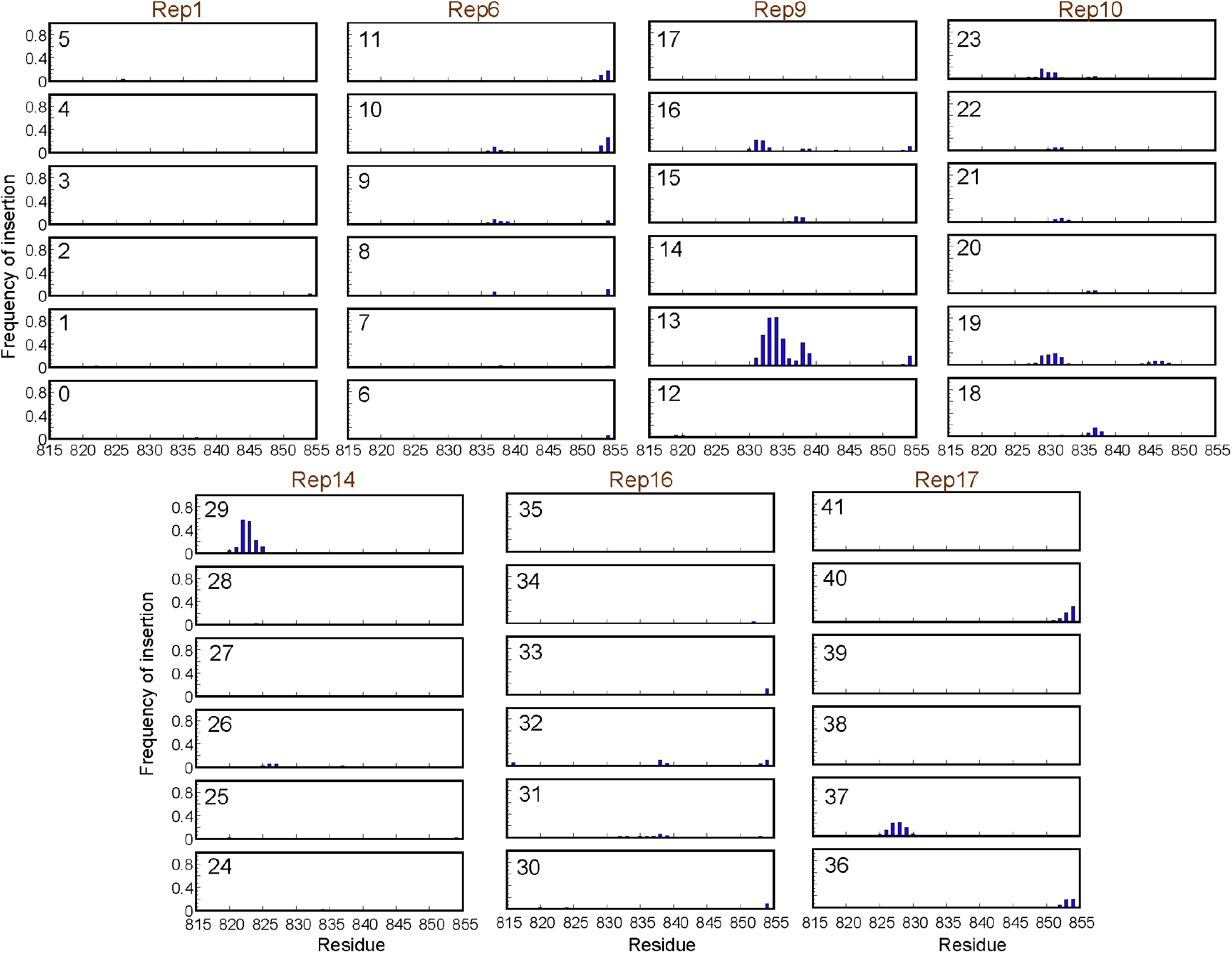
Frequency of membrane insertion for each residue of the SARS-CoV-2-FP plotted separately for each MD trajectory calculated the peptide-membrane system. Each column presents data for the set of simulations initiated from the same conformation and labeled according to the scheme used in the main text, Figures 2 and 3. A residue was considered inserted into the membrane if the z distance between its Cα atom and the C22 backbone carbons was <5Å.

**Figure S7:**
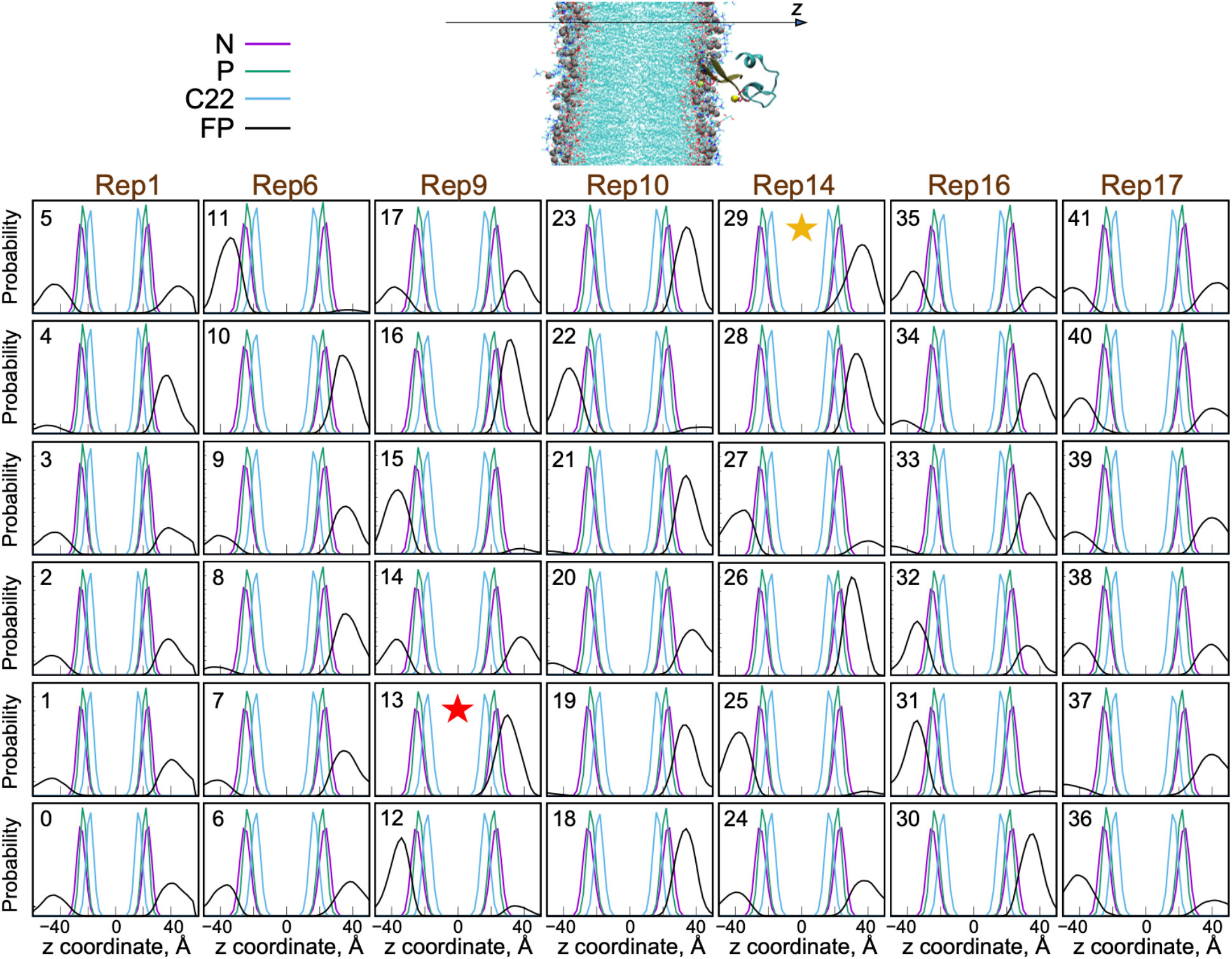
The grid of panels presents the probability of finding a POPC lipid head-group nitrogen atom (N), phosphorus atom (P), hydrocarbon chain C22 carbon (C22), any atom of the SARS-CoV-2-FP (FP), and the F833/I834 residues of the SARS-CoV-2-FP (F833/I834) in 2Å rectangular slabs along membrane-normal z axis in all the MD simulations of the SARS-CoV-2-FP-membrane system. Each column presents data for the set of simulations initiated from the same conformation and labeled according to the scheme used in the main text, Figures 2 and 3. The red and yellow stars highlight the trajectories 13 and 29 in Rep9 and Rep14, respectively, in which the largest extent of the peptide insertion into the membrane was observed and which are described in detail in Figure 4 of the main text. The molecular representation of the system shown above (middle) depicts the peptide-bound bilayer in the orientation that matches the direction of the x-axis in the panels below it.

**Figure S8:**
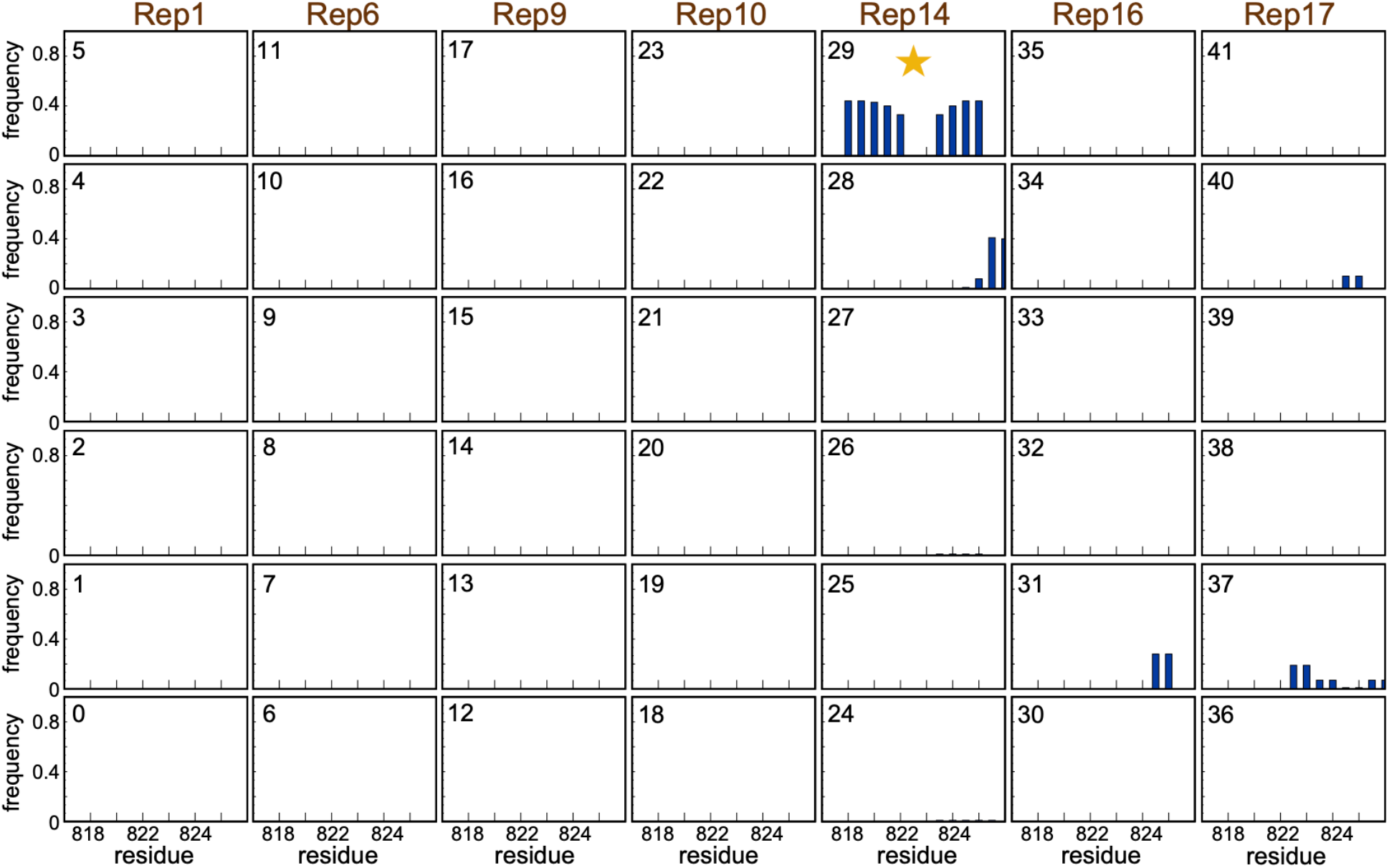
Frequency of finding the N-terminal residues (816-826) in the extended, beta-sheet conformation in the simulations of the SARS-CoV2-FP-membrane systems. The N-terminal segment assumes this extended conformation when the peptide establishes Mode 2 membrane binding (Rep14, highlighted with a yellow star).

**Figure S9:**
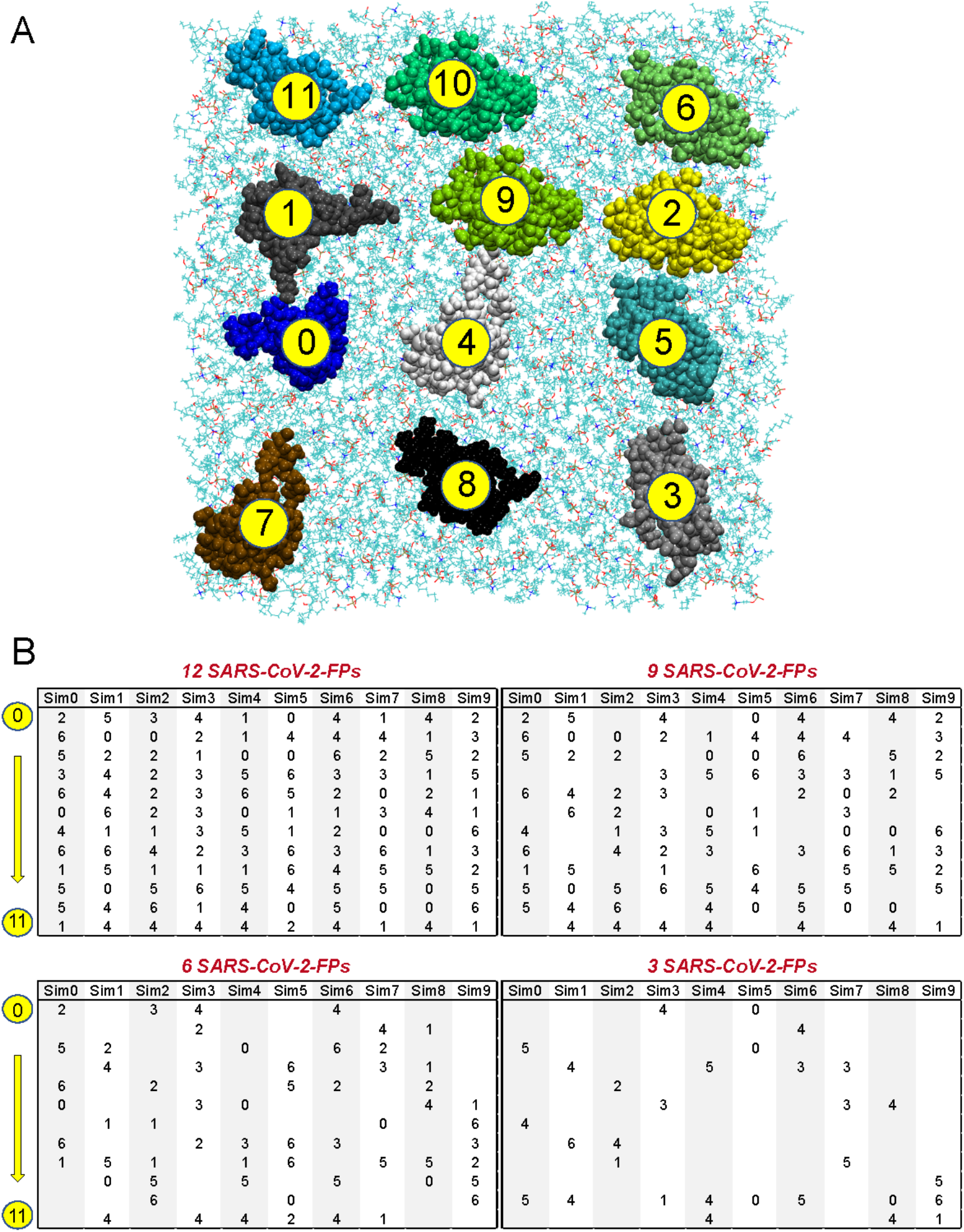
(**A**) A snapshot illustrating SARS-CoV-2-FPs on a 3×4 grid (positions labeled from 0 to 11) next to the 3:1:1: POPC/POPG/Chol membrane. The 12 FPs are shown in space fill, and the membrane components are rendered in lines. (**B**) The protocol for random placement of 12, 9, 6, or 3 SARS-CoV-2-FPs on the grid shown in panel A. Starting with the seven Ca^2+^-bound conformations of SARS-CoV-2-FP (from Figure 2B), 12 conformations were randomly selected (thus models appear more than once in a selection), and placed on the grid as shown in **(A)**(0⇒11). The 7 conformations from Figure 2B are labeled from 0 to 6 using the following convention: Rep1 – 0, Rep6 – 1, Rep9 – 2, Rep10 – 3, Rep14 – 4, Rep16 – 5, Rep17 – 6. This To create 10 different starting grids of 12 FP structures (Sim0 to Sim9), this procedure was repeated 10 times. To create systems with 9, 6, and 3 FPs, we randomly removed an appropriate number of FPs from the grids of 12 FPs.

**Figure S10:**
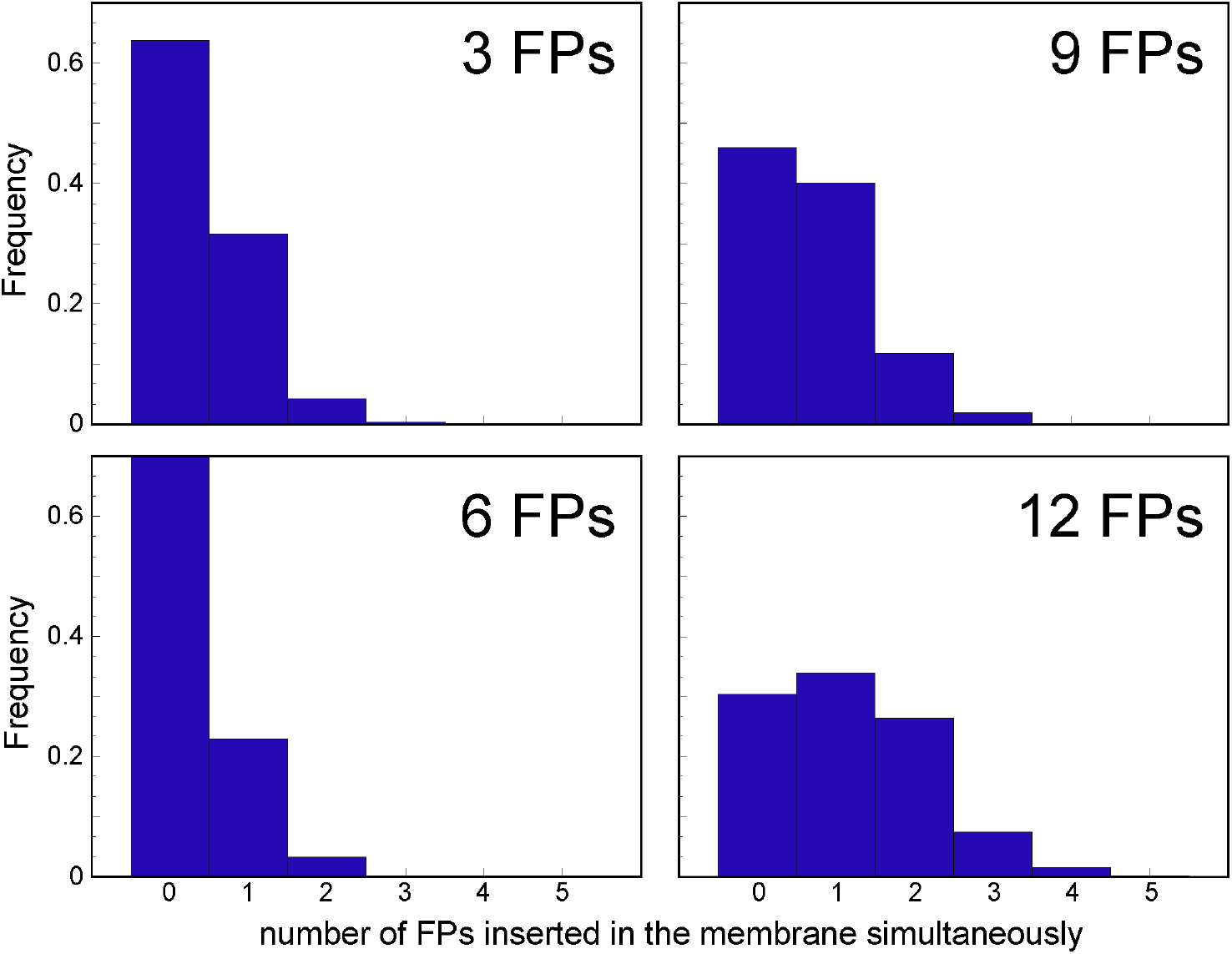
Fraction of trajectory frames (y axis) with a specific number (x axis) of FPs simultaneously inserted into the membrane in the simulations of 3, 6, 9, and 12 SARS-CoV-2-FP systems.

**Figure S11:**
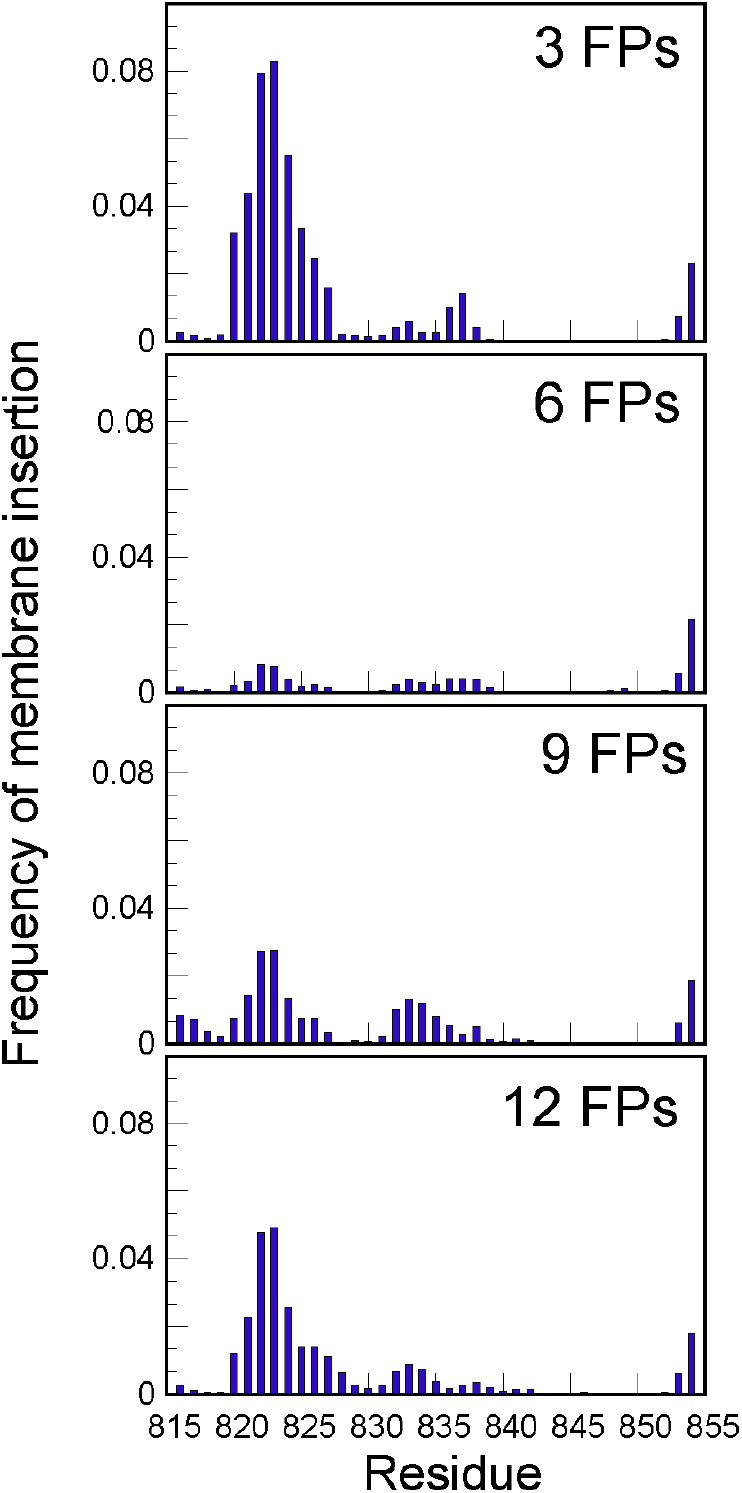
Frequency of membrane insertion for each residue of SARS-CoV-2-FP from analysis of the MD simulations of 3, 6, 9, and 12 FPs interacting with 3:1:1 POPC/POPG/Cholesterol bilayer. The data shown are the average over 60 trajectories per each system (see Table 1). Membrane insertion was defined as described in Figure 3B caption.

**Figure S12:**
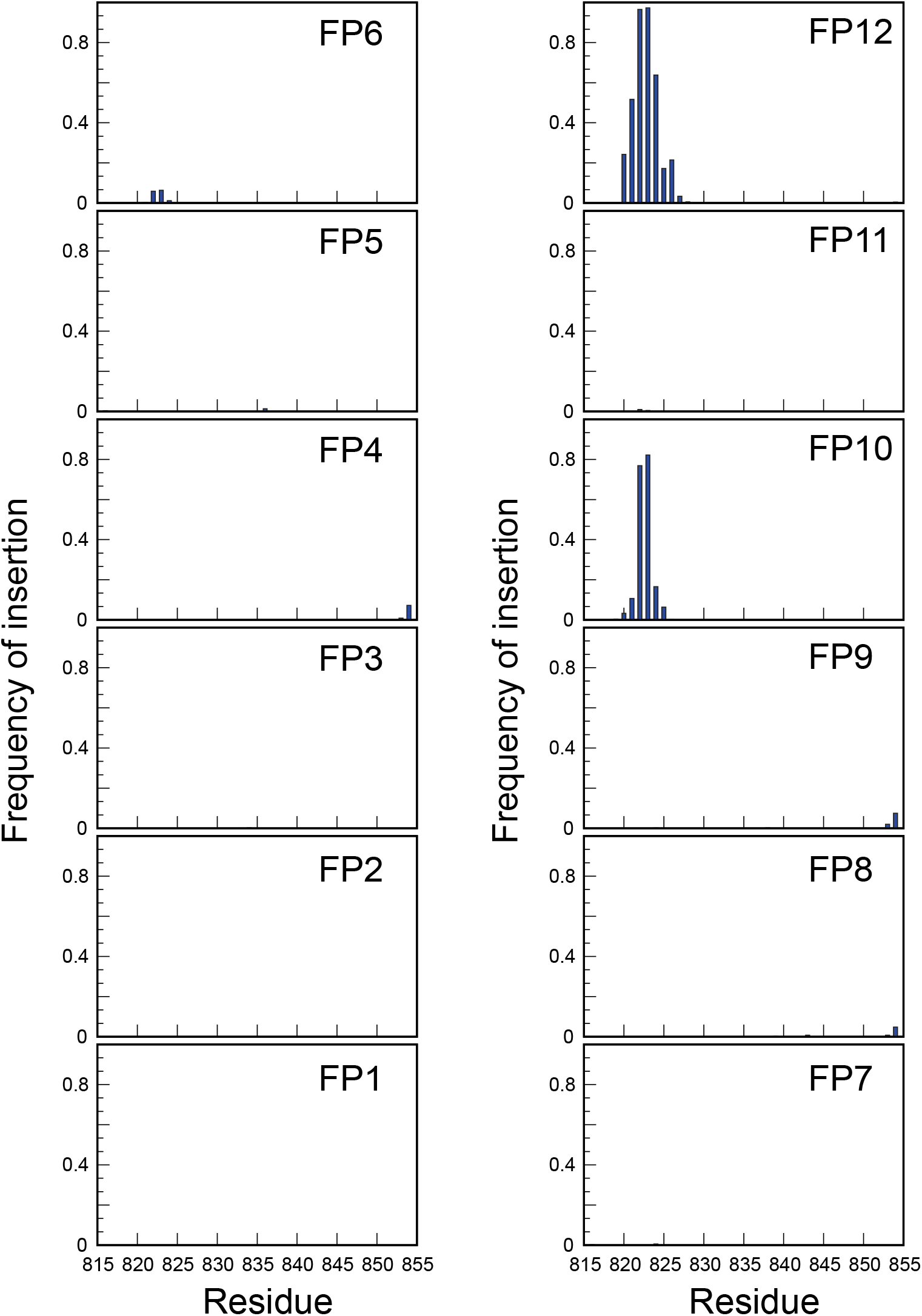
Frequency of membrane insertion for each residue of SARS-CoV-2-FP from analysis of the MD simulation of the 12 FPs construct interacting with 3:1:1 POPC/POPG/Cholesterol bilayer for which the pressure profile analysis was carried out. The different panels show data separately for different FPs in the system. Membrane insertion was defined as described in Figure 3B caption.

**Figure S13:**
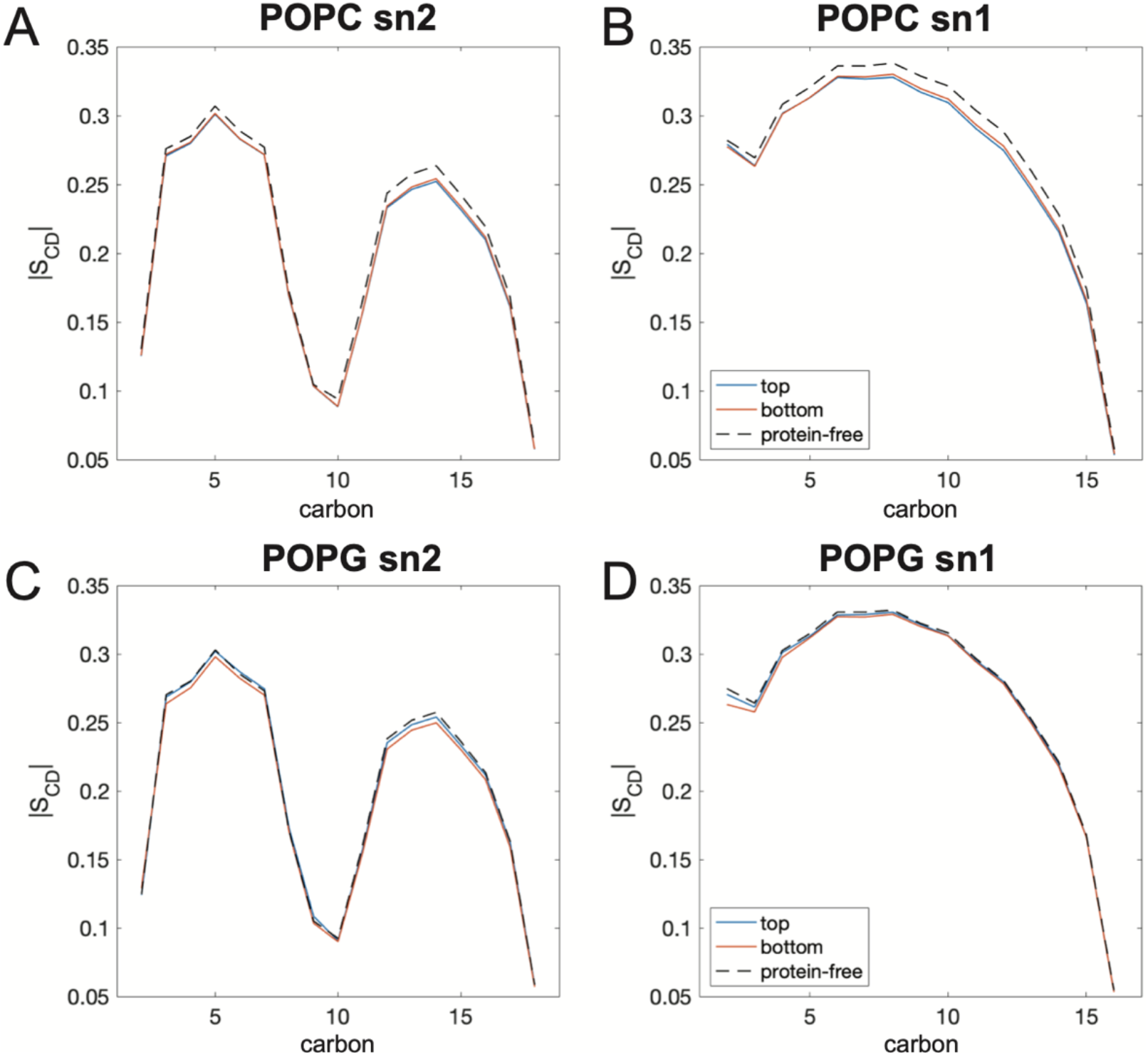
Deuterium order parameters for carbon atoms in the sn2 (A,C) and sn1 (B,D) hydrocarbon chains of POPC (A-B) and POPG (C-D) lipids calculated in the two leaflets of the membrane (*top*, *bottom*) from the simulations of 12FP-membrane system (+12FP) used for the analysis of the pressure profile. For reference, order parameters of the protein-free system are also shown in dashed lines.

**Figure S14:**
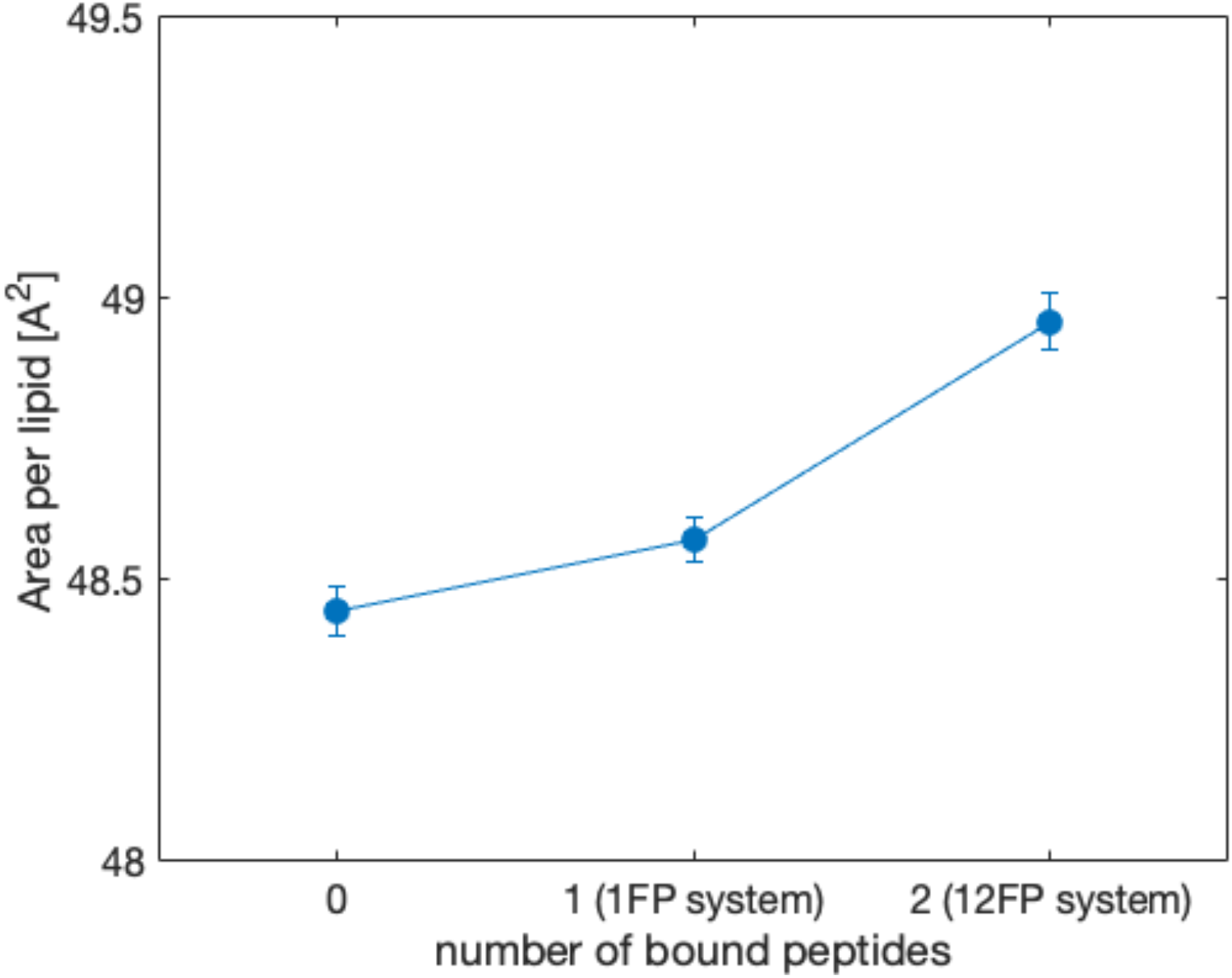
Area per lipid as a function of the number of bound peptides as quantified from the following simulations: one of the 1FP MD simulations from the Rep0 set that has no FP insertion(i.e., simulation ID 0 in Figure 3C) shown as 0 on the x-axis (number of bound peptides); one of the 1FP MD simulations from the Rep14 set with one FP inserted in the membrane (i.e., simulation ID 29 in Figure 3C) indicated as 1FP system on the x-axis, and one of the MD simulations for the 12FP system with 2FPs inserted in the membrane (number of bound peptides: 2) indicated as 12FP system on the x-axis. The error bars were estimated from bootstrapping analysis of the corresponding trajectories.

